# LKOM: Longitudinal Knockoff with Masked Likelihood Ratio for FDR-Controlled Association Discovery

**DOI:** 10.64898/2026.10.15.759368

**Authors:** Yundi Hu, He Xu, Shiyang Ma, Wenjian Bi, Xiaolei Lin

## Abstract

Repeated phenotypic measurements in biobanks can reveal genetic effects on both average trait levels and within-subject variability. However, existing longitudinal genome-wide association methods generally test variants marginally and analyze the two components separately, limiting the localization of signals in linkage disequilibrium and the detection of variants with shared effects. Here we introduce Longitudinal KnockOff with Masked likelihood ratio (LKOM), a multivariable framework that jointly models mean and variability signals while controlling the variant-level false discovery rate for the joint null of no association with either component. LKOM uses model-X knockoffs and masked likelihood-ratio statistics to distinguish original variants from synthetic controls, and assigns selected variants to mean-specific, variability-specific or shared association states. A genetic-relationship-matrix-based whitening extension accommodates related individuals. In simulations, LKOM maintained false discovery rate control across diverse genetic architectures, heavy-tailed errors, reduced longitudinal follow-up and sample sizes up to 100,000. It generally improved power over separate-channel knockoff and marginal association methods, with the largest gains for shared effects and rare variants, particularly when the overall genetic contribution to phenotypic variation was modest. In related samples, whitening restored calibration under dense pedigree structures while preserving power gains over SPA_**GRM(CCT)**_. LKOM provides an interpretable and scalable approach for identifying conditionally associated variants from longitudinal phenotypes.

## Introduction

Large-scale biobanks and electronic health records increasingly provide repeated phenotypic measurements alongside genomic data [1, 2]. Compared with cross-sectional phenotypes, longitudinal phenotypes capture dynamic features such as disease progression, changes in metabolic status, and within-individual variability [2]. For example, the long-term level and variability of glucose, blood pressure, as well as body mass index are all related to disease risk, highlighting the potential relevance of variability beyond average exposure [3–5]. Therefore, analyses based on one observation or an individual-level average may miss genetic effects on within-subject (WS) variability. Longitudinal location-scale models account for dependence among repeated measurements while allowing predictors to affect both the mean and WS variance [6]. Building on this framework, TrajGWAS brought this approach to biobank-scale association testing [2]. More recently, SPA_GRM_ extended analysis to related individuals and improved the calibration of association tests across challenging genotype and phenotype distributions [7]. These developments have established the feasibility of large-scale genetic analyses of longitudinal mean and variance effects.

Most longitudinal genome-wide association analyses test variants individually. Although effective for discovering associated regions, marginal tests have limited ability to distinguish signals from correlated variants within regions of linkage disequilibrium (LD) [8, 9]. A variant may be marginally associated with a phenotype because it tags another associated variant. Mistaking such a proxy for the variant contributing independent association evidence can misdirect functional follow-up and biological interpretation [10, 11]. Conditional and joint association analyses address this problem by evaluating multiple variants together [12], while Bayesian fine-mapping methods, including the sum of single effects model, quantify uncertainty over alternative configurations of associated variants [13]. These approaches motivate a multivariable perspective in which variants are evaluated conditional on other variants included in the analysis.

A discovery procedure must also address errors among the variants it selects. The false discovery rate (FDR) is the expected proportion of false discoveries in a selected set [14]. For multivariable selection, the target is whether a variant provides additional information about the phenotype conditional on the other analyzed variants and covariates [15]. Multiple-testing correction of marginal associations does not address this conditional target, and multivariable modeling alone does not automatically provide frequentist FDR control. The knockoff framework offers a principled approach to conditional variable selection by introducing synthetic controls against which the original variants compete [16]. In the model-X formulation, pairwise exchangeability and conditional independence are the key conditions underlying knockoff FDR control [15]. Genomic applications using hidden Markov model knockoffs and Knockoff-Screen demonstrate the utility of this principle for discovery in the presence of LD [17, 18]. Nevertheless, the power of a knockoff procedure depends on how effectively its feature statistics distinguish original variables from their synthetic counterparts. Masked likelihood ratio (MLR) statistics address this distinction directly by assessing evidence about the identities of original-knockoff pairs under a working Bayesian model [19]. This provides a principled starting point for constructing statistics that use the complementary information contained in longitudinal mean and variance effects.

A further challenge is that the two longitudinal components need not be equally informative. Variance-effect inference can be sensitive to asymmetric residual distributions and limited information about individual variability [2, 6]. These can leave weak variance associations undetected when each channel is analyzed separately. In an analysis of 79 longitudinal traits, SPA_GRM_ reported that 313 of 362 loci associated with WS variability also showed associations with mean levels [7], which suggests that the two components can share genetic signals. This motivates a model that allows shared as well as component-specific genetic effects. More generally, joint modeling of related genetic effects can improve inference by learning and exploiting patterns of sharing [20]. In the longitudinal setting, a variant may affect neither component, the mean only, the variance only, or both. A model that accommodates these four possibilities can borrow information when the data support sharing without requiring every association to occur in both channels. However, no existing knockoff framework jointly uses the mean and WS variability components for FDR-controlled variant selection while also characterizing component-specific and shared associations.

Here, we introduce LKOM, a Longitudinal KnockOff with Masked likelihood ratio with three main methodological contributions. First, LKOM constructs dual-channel masked likelihood ratio statistics that jointly use evidence from the mean and WS variability components to select variants against the joint null of no effect on either component. Second, its Bayesian working model provides posterior probabilities for the four association states: null, mean only, WS variability only, and shared effects. Third, a GRM-based extension accommodates related individuals while preserving symmetry between original and knockoff genotypes. Across extensive simulations, LKOM controlled the FDR and achieved higher power than separate-channel knockoff and marginal association methods, with the largest gains when signals were shared between the two channels.

## Results

### Overview of the LKOM framework

LKOM identifies genetic variants associated with the mean or within-subject (WS) variability of longitudinal traits while controlling the false discovery rate (FDR) at the variant level. The framework consists of four steps (Fig. 1). First, we fit a covariate-adjusted null model using WiSER and derive subject-level residual scores for the mean and WS-variability channels. A rank-based inverse-normal transformation (INT) is applied separately to each channel before joint modeling.

**Fig. 1.**
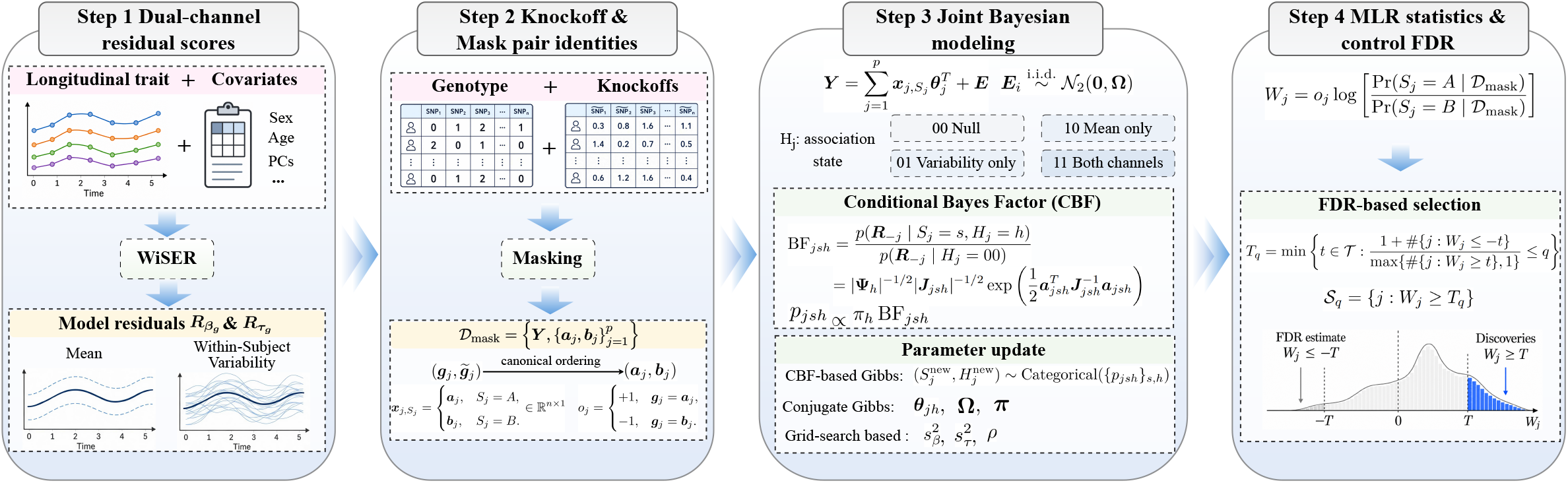
Overview of the LKOM framework. **Step 1**, A WiSER null model adjusts longitudinal phenotypes for covariates, including sex, age and genetic principal components (PCs), and produces subject-level residual scores for the mean 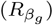 and within-subject (WS) variability 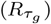. The two channels undergo separate rank-based inverse-normal transformations to form the response matrix **Y. Step 2**, Knockoff genotypes are generated from the original genotype matrix. Each original–knockoff pair (*g*_*j*_, 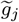) is canonically ordered as (*a*_*j*_, *b*_*j*_ ), yielding masked data *D*_mask_ that conceal the original-versusknockoff identities. The latent variable *S* specifies which member is the original genotype; *o* records its observed position in the canonical pair. **Step 3**, A joint Bayesian regression models the two channels, with ***θ***_*j*_ denoting genetic effects and **Ω** the residual covariance matrix. The association state *H*_*j*_ distinguishes null (00), mean-only (10), WS-variability-only (01) and shared (11) effects. Conditional Bayes factors (CBFs) support joint updates of pair identities and association states, alongside updates of model parameters. **Step 4**, Posterior pair-identity odds and the observed orientation *o*_*j*_ yield the masked likelihood ratio (MLR) statistic *W*_*j*_ . The knockoff+ filter compares the numbers of statistics below *−t* and above *t* to determine the threshold *T*_*q*_ at target false discovery rate (FDR) *q*. Variants with *W*_*j*_ *≥ T*_*q*_ form the discovery set *S*_*q*_.

Second, we generate knockoff genotypes as negative controls that mimic the correlation structure of the original genotypes. A symmetric marginal screen retains promising variant–knockoff pairs for subsequent analysis (Methods). Each pair is then placed in a canonical order that does not depend on which member is the original genotype, masking the original-versus-knockoff identities from the model. Third, we jointly model the two residual channels using a Bayesian regression with four association states: null (00), mean only (10), WS variability only (01), and both channels (11). Conditional Bayes factors are used in a Gibbs sampler to infer the masked pair identities and association states while accounting for the effects of other modeled variants. This formulation accommodates both shared and channel-specific genetic effects.

Finally, we construct a masked likelihood ratio (MLR) statistic from the posterior odds favoring the original genotype over its knockoff. The statistic changes sign when the original and knockoff labels are exchanged. The knockoff+ filter uses large negative statistics to estimate the number of false discoveries among variants with large positive statistics and selects variants at the target FDR. Posterior association-state probabilities additionally indicate whether the evidence for each selected variant arises from the mean channel, the WS-variability channel, or both. For cohorts containing related participants, a GRM-based whitening step is applied to the residual channels and both genotype matrices before joint modeling.

### LKOM maintains well-controlled FDR across diverse genetic architectures

We conducted simulation studies using UK Biobank-derived genotypes from 10,000 participants to evaluate the FDR control and power of LKOM for detecting variants associated with the trajectory mean, WS variability, or both. FDR and power were summarized across 500 evaluations per setting (Methods). FDR was evaluated at the variant level relative to the joint null hypothesis *H*_0,*j*_ : *β*_*j*_ = *τ*_*j*_ = 0. Because LKOM jointly models the 2 channels, we first examined whether genetic signals in 1 channel interfered with inference in the other and inflated the joint FDR. We considered 4 association states: the global null, mean effects only, WS variability effects only, and effects on both channels. The 3 nonnull states were evaluated under low, medium, and high proportions of variance explained (PVE), with 20 causal variants in each active channel. The low, medium, and high PVE settings were (PVE_*β*_, PVE_*τ*_ ) = (0.025, 0.05), (0.05, 0.10), and (0.10, 0.20). Given the greater difficulty of detecting genetic effects on WS variability, we assigned a higher PVE to this channel. Under the global null, LKOM made no discoveries at any target FDR level examined, yielding an observed FDR of 0. Across the 3 nonnull states, the observed FDR ranged from 2.34% to 5.67% at a target level of 5%, and from 7.38% to 11.49% at a target level of 10% (Fig. 2A). When both channels were affected, the observed FDR remained close to the target level. Modest upward deviations occurred at high PVE, where the observed FDR reached 5.67% and 11.49% at target levels of 5% and 10%, respectively.

**Fig. 2.**
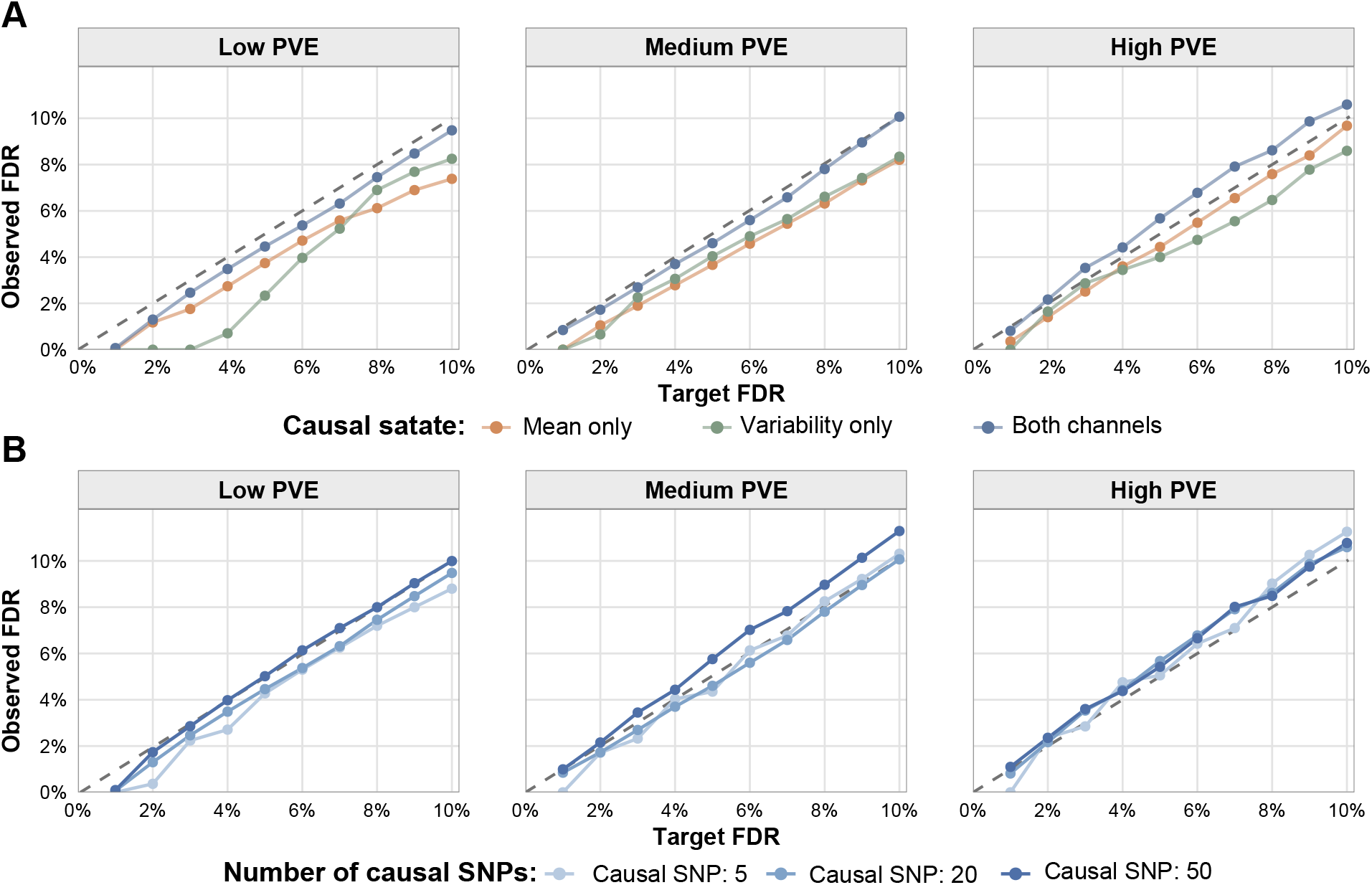
False discovery rate control of LKOM across genetic architectures. **A**, Observed false discovery rate (FDR) under mean-only, within-subject (WS) variability-only, and two-channel genetic effects, with 20 causal SNPs in each active channel. **B**, Observed FDR with 5, 20, or 50 causal SNPs per channel. Low, medium, and high proportions of variance explained (PVE) correspond to (PVE_*β*_, PVE_*τ*_ ) = (0.025, 0.05), (0.05, 0.10), and (0.10, 0.20), respectively. Simulations included 10,000 participants with 6–15 longitudinal measurements each. FDR was evaluated at the variant level relative to the joint null hypothesis *β*_*j*_ = *τ*_*j*_ = 0. Points show the mean false discovery proportion across 500 grouped evaluations, each pooling SNP-level results from 5 independent simulation replicates. Grey dashed lines indicate equality between observed and target FDR.

We next examined whether FDR control depended on signal sparsity. We varied the number of causal variants per channel over 5, 20, and 50 and considered the same 3 PVE levels. At each PVE level, increasing the number of causal variants while keeping the total genetic contribution within each channel fixed reduced the average effect size per variant. Across the 9 combinations of causal variant number and PVE, the observed FDR ranged from 4.27% to 5.76% at a target level of 5%, and from 8.80% to 11.49% at a target level of 10% (Fig. 2B).

We further evaluated asymmetric genetic architectures to determine whether differences between the 2 channels could produce excessive false discoveries. To vary signal number, we fixed 1 channel at 20 causal variants and varied the other over 5, 10, and 15. Both directions of imbalance were considered. We also considered settings with unequal PVE between the 2 channels. When the numbers of causal variants differed, the observed FDR ranged from 4.51% to 5.62% at the 5% target and from 9.65% to 10.89% at the 10% target. Under unequal PVE, the corresponding ranges were 5.17% to 5.75% and 10.03% to 11.00%, respectively (Supplementary Fig. S1). Overall, LKOM maintained observed FDR close to the nominal levels across different association states, numbers of causal variants, PVE levels, and imbalances in signal number or strength between the 2 channels.

### LKOM improves power across diverse genetic architectures

We next evaluated whether LKOM improved the detection of causal variants. We compared LKOM with LKOM-single, TrajGWAS, and TrajKnock. LKOM-single applied the same MLR framework separately to the 2 response channels, thereby isolating the contribution of joint modeling. TrajKnock applied TrajGWAS separately to the original and knockoff genotypes within each channel, providing a benchmark for knockoff-based selection without MLR and joint modeling. For all 4 methods, power was calculated separately for each channel and then averaged across the 2 channels. For LKOM, mean-channel power was the proportion of truly beta-causal SNPs that were selected and assigned to state 10 or 11, same for WS variability-channel power. The knockoff-based methods were evaluated at a target FDR of 5%. For TrajGWAS, which tests marginal SNP associations, we used a Bonferroni-corrected significance level of 0.05.

Power increased with PVE and decreased as genetic effects were distributed across more causal variants at a given PVE level (Fig. 3A). Across the 9 causal-number and PVE combinations, LKOM outperformed TrajGWAS in 8 settings, with power gains ranging from 4.72% to 16.73%. The only exception was the most diffuse low-PVE setting with 50 causal variants per channel, where the power of LKOM and TrajGWAS was 17.48% and 18.52%, respectively. LKOM consistently outperformed LKOM-single, although the gains were generally smaller, ranging from 0.35% to 16.28%. Compared with TrajKnock, LKOM increased power by 3.48% to 51.18%, and TrajKnock had the lowest power in all 9 settings.

**Fig. 3.**
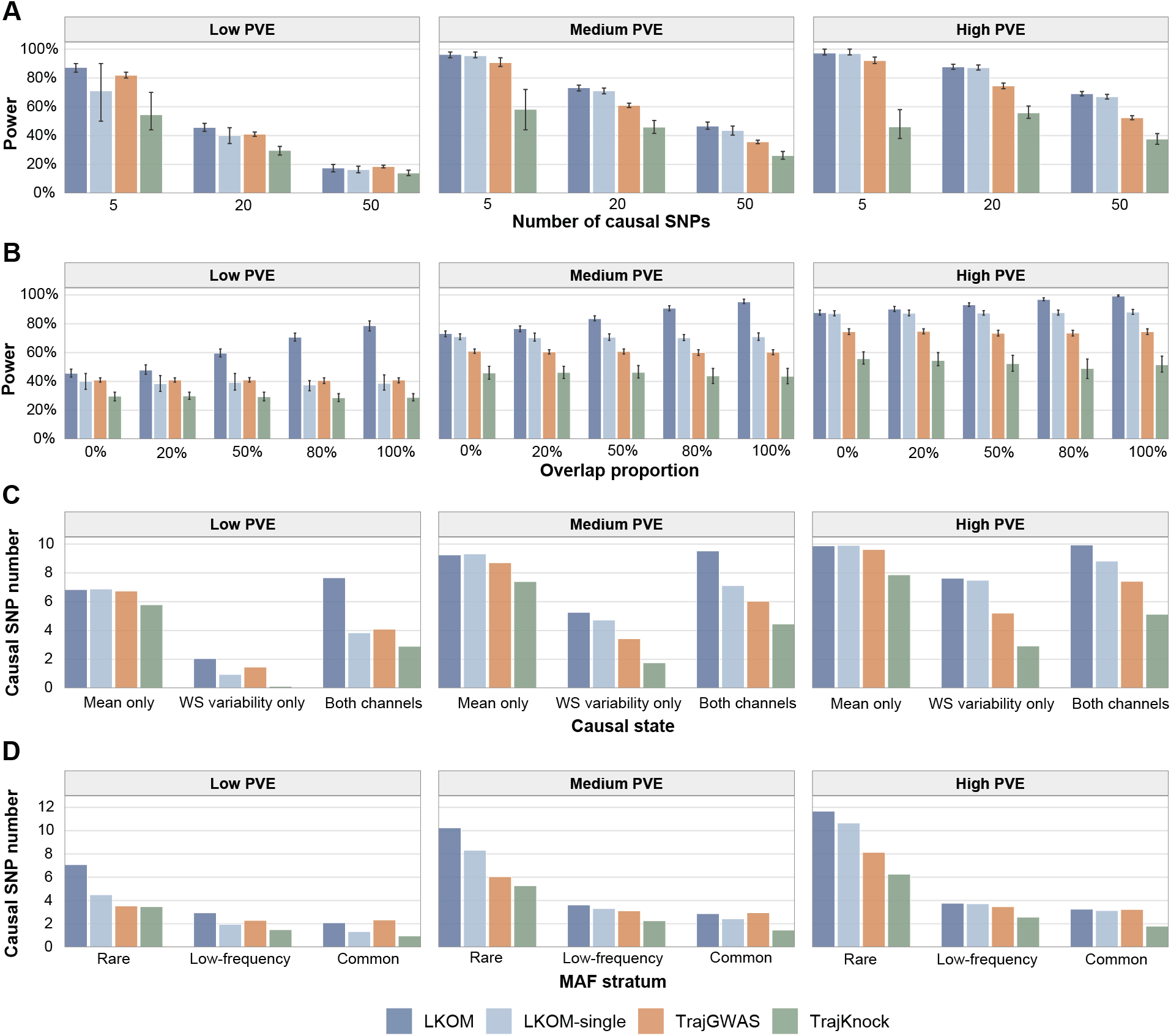
Power comparison and sources of detection gains in LKOM. **A**, Power with 5, 20, or 50 causal SNPs per channel. **B**, Power across overlap proportions of 0, 0.2, 0.5, 0.8, and 1 between the 2 causal sets. **C**, Mean numbers of detected causal SNPs stratified by their true association state: mean only, within-subject (WS) variability only, or shared effects. Detections are counted within the affected channel for channel-specific variants and averaged across the 2 channels for shared-effect variants. **D**, Mean numbers of detected causal SNPs per channel within rare (1.25 *×* 10^*−*3^ *≤* MAF *<* 0.01), low-frequency (0.01 *≤* MAF *<* 0.05), and common (0.05 *≤* MAF *≤* 0.5) strata, where MAF denotes minor allele frequency. Panels **B–D** used 20 causal SNPs per channel; counts in **C** and **D** are reported per simulation replicate and averaged across the 5 overlap settings. Low, medium, and high proportions of variance explained (PVE) correspond to (PVE_*β*_, PVE_*τ*_ ) = (0.025, 0.05), (0.05, 0.10), and (0.10, 0.20), respectively. Simulations included 10,000 participants with 6– 15 longitudinal measurements each. This power is the average of the proportions of true causal SNPs detected in the mean and WS variability channels. For LKOM, a true beta-causal SNP is counted when selected and assigned to state 10 or 11; a true tau-causal SNP is counted when selected and assigned to state 01 or 11. LKOM-single models each channel separately, whereas TrajKnock applies knockoff selection using channel-specific TrajGWAS statistics. The knockoff-based methods used a target false discovery rate of 5%; TrajGWAS used a Bonferroni-corrected significance level of 0.05. Bars show means across 500 grouped evaluations, each pooling 5 replicates sampled from 200 independent simulations per setting. Error bars in **A** and **B** indicate the 25th and 75th percentiles.

LKOM also retained its power advantage under asymmetric genetic architectures. When the numbers of causal variants differed between the 2 channels, LKOM achieved the highest power in all 6 settings and exceeded LKOM-single by 2.58% to 4.05%. When the channel-specific PVE values differed, LKOM again had the highest power in all 6 settings, with gains of 0.44% to 6.80% over LKOM-single (Supplementary Fig. S2, S3).

Substantial sharing of genetic effects between the mean and WS variability components has been observed in longitudinal trait analyses. Across 79 longitudinal traits in UK Biobank, Xu et al. reported that 313 of 362 loci associated with WS variability also influenced mean levels [7]. We therefore varied the overlap proportion between the 2 causal sets over 0, 0.2, 0.5, 0.8, and 1. The power of LKOM increased markedly with the degree of overlap, whereas the power of the 3 separate-channel methods remained largely unchanged (Fig. 3B). Averaged across the 3 PVE levels, the gains of LKOM over TrajGWAS were 10.06%, 12.83%, 20.38%, 28.03%, and 32.45% at corresponding overlap proportion, respectively. The corresponding gains over LKOM-single were 2.80%, 6.20%, 12.99%, 20.88%, and 25.11%. The advantage over LKOM-single was also greater under weaker signals, averaging 21.72%, 13.19%, and 5.88% across the 5 overlap proportions under low, medium, and high PVE, respectively. In comparison, the average gains over TrajGWAS were 19.57%, 23.37%, and 19.31% across the same PVE levels. These gains increased with cross-channel sharing, with the largest improvement over LKOM-single occurring under low PVE.

Because averaging power across the 2 channels does not require recovery of the complete association state, we additionally evaluated exact-state power, defined as the proportion of causal variants that were both selected and assigned to their true state among 10, 01, and 11. Exact-state power showed a similar dependence on cross-channel sharing, with particularly large gains at high overlap. Averaged across the 3 PVE levels, the gains over TrajGWAS were 10.32%, 11.72%, 20.58%, 33.36%, and 57.66% across increasing overlap levels, compared with 2.60%, 4.42%, 11.62%, 24.28%, and 47.27% over LKOM-single. Across overlap levels, the gains over LKOM-single averaged 29.48%, 18.08%, and 6.55% under low, medium, and high PVE, respectively, while the corresponding gains over TrajGWAS were 24.97%, 29.79%, and 25.43% (Supplementary Fig. S5). The confusion matrices showed that most detected variants were assigned to their correct association states. Misclassification was most apparent for shared 11 variants at low PVE, which were more often assigned to state 10 than to state 01. These errors diminished as the signal strength increased (Supplementary Fig. S6).

### Power gains are concentrated among shared-effect and rare variants

We next examined which causal states contributed to the detection advantage of LKOM after pooling results across the five overlap settings (Fig. 3C). The advantage of LKOM was most pronounced for the both-channels state. Relative to LKOM-single, LKOM detected an average of 3.82, 2.41, and 1.12 more causal variants per channel per simulation replicate under low, medium, and high PVE, respectively. The corresponding gains over TrajGWAS were 3.57, 3.49, and 2.52 variants. LKOM also improved the detection of variants affecting WS variability only, with gains of 1.10, 0.54, 0.14 variants over LKOM-single and 0.59, 1.83, 2.42 variants over TrajGWAS. By contrast, for the mean-only state, LKOM detected slightly fewer causal variants than LKOM-single, although the differences were minimal at *−*0.04, *−*0.07, and *−*0.03 variants across the three PVE levels. The overall power improvement of LKOM was therefore driven primarily by its improved detection of variants affecting both the mean and WS variability components. For these shared-effect variants, the advantage of joint modeling became more pronounced as PVE decreased.

We also examined which MAF strata contributed to the detection advantage of LKOM. We also compared the mean numbers of detected causal variants, averaged across the 5 overlap settings (Fig. 3D). Variants were classified as rare (MAF *<* 0.01), low-frequency (0.01 *≤* MAF *<* 0.05), or common (MAF *≥* 0.05). LKOM detected more causal variants than LKOM-single across all 3 MAF strata, with the largest improvement observed for rare variants. Under low, medium, and high PVE, LKOM detected 2.58, 1.92, and 1.02 more rare causal variants per channel per simulation replicate than LKOM-single, respectively. The corresponding gains were 0.99, 0.31, and 0.04 for low-frequency variants and 0.75, 0.44, and 0.11 for common variants. Compared with TrajGWAS, LKOM detected 3.54, 4.21, and 3.54 additional rare variants and 0.65, 0.50, and 0.30 additional low-frequency variants across the three PVE levels. By contrast, the two methods detected similar numbers of common causal variants, with 2.08, 2.86, 3.25 for LKOM versus 2.33, 2.95, 3.23 for TrajGWAS under low, medium, and high PVE. The additional detections by LKOM were therefore concentrated among rare variants.

### LKOM maintains robust performance in related samples through GRM-based whitening

Sample relatedness is common in large biobank cohorts and can inflate false-positive rates when the correlation between related individuals is ignored. Restricting analyses to unrelated participants avoids this problem but discards potentially informative samples and consequently reduces statistical power [7]. To accommodate related individuals, we extended LKOM by applying GRM-based whitening to its channel-specific working responses and the corresponding original and knockoff genotype matrices. This transformation accounts for the modeled covariance between participants before construction of the knockoff statistics. We compared LKOM with SPA_GRM_, a recently developed method for longitudinal-trait GWAS with related samples [7], using its Cauchy combination test implementation, SPA_GRM(CCT)_. This implementation showed consistently high power in the simulations reported in the original study.

We first simulated regular pedigrees containing 4, 10, or 20 members using the GRAB package and UK Biobank genotype data (Supplementary Method S10). The resulting GRMs showed clearly delineated blocks of related individuals corresponding to the pedigrees (Fig. 4A). Without whitening, the observed FDR exceeded the nominal level, reaching 11.98%, 12.05%, and 12.32% at a target FDR of 10% for the 4-, 10-, and 20-member pedigrees, respectively. Whitening reduced the corresponding values to 6.42%, 6.10%, and 7.50%(Fig. 4B). Although the observed FDR was somewhat conservative after whitening, this was accompanied by only a modest loss of power, with reductions of 2.39%, 2.96%, and 2.81% for the 4-, 10-, and 20-member pedigrees, respectively. Nevertheless, whitened LKOM remained substantially more powerful than SPA_GRM(CCT)_, exceeding it by 14.71%, 15.20%, and 16.03%, respectively (Fig. 4C).

**Fig. 4.**
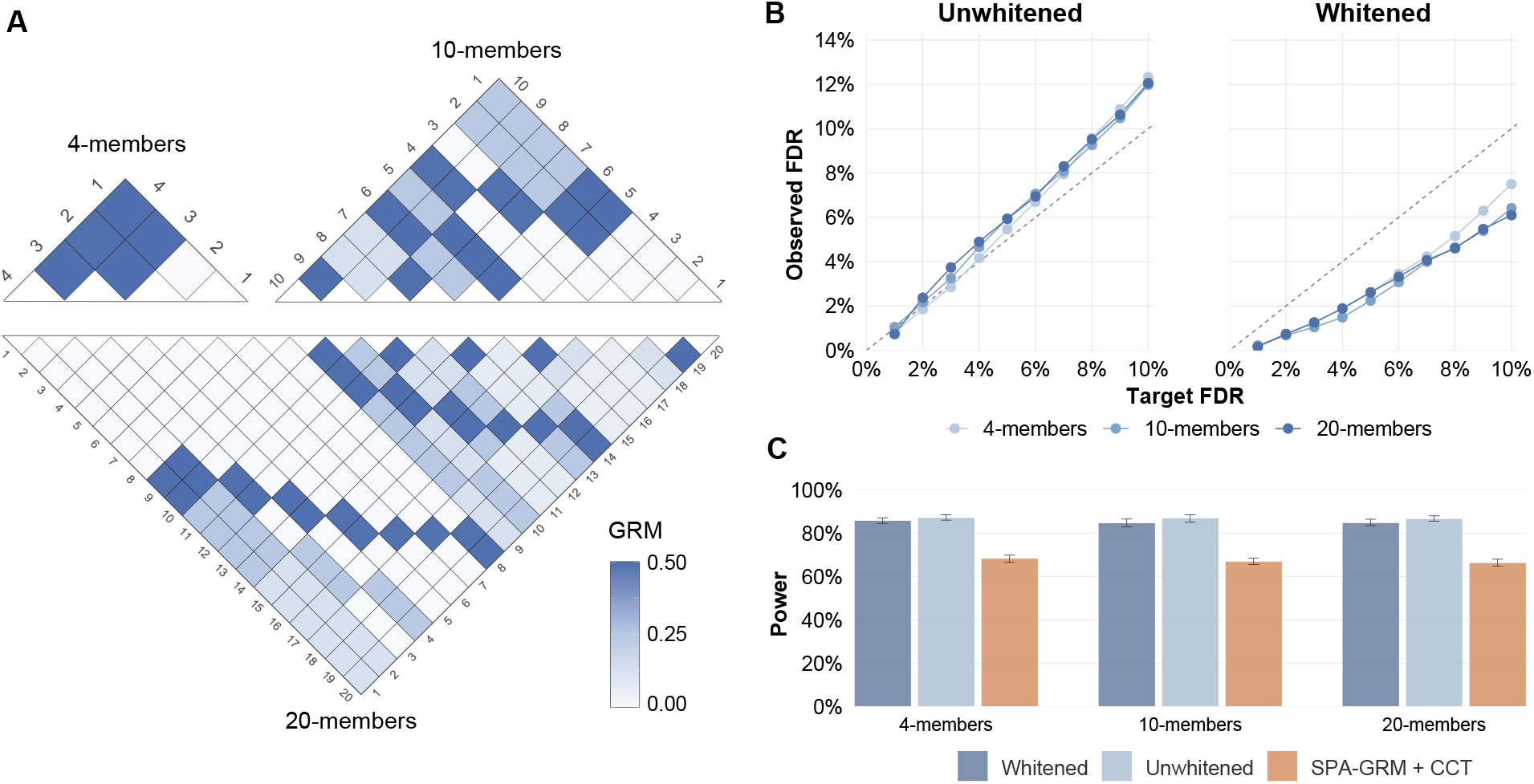
Effect of GRM-based whitening on LKOM in related samples. **A**, Within-pedigree genetic relationship matrices (GRMs) for simulated families containing 4, 10, or 20 members. Colour intensity indicates pairwise GRM entries. **B**, Observed false discovery rate (FDR) of LKOM before and after GRM-based whitening across the 3 family sizes. Grey dashed lines indicate equality between observed and target FDR. **C**, Channel-wise power of whitened LKOM, unwhitened LKOM, and SPA-GRM with the Cauchy combination test (CCT). Pedigrees were generated using the GRAB package and UK Biobank genotype data. Simulations included 10,000 participants with 6–15 longitudinal measurements each, 20 causal SNPs per channel, and proportions of variance explained of 0.05 for the mean channel and 0.10 for the within-subject (WS) variability channel. Whitening was applied to the channel-specific working responses and the corresponding original and knockoff genotype matrices. FDR was evaluated against the joint null hypothesis *β*_*j*_ = *τ*_*j*_ = 0, and channel-wise power was averaged across the mean and WS variability channels. In **C**, LKOM used a target FDR of 5%, and SPA-GRM+CCT used a Bonferroni-corrected significance level of 0.05. Points and bars show means across 500 grouped evaluations per setting. Error bars in **C** indicate the 25th and 75th percentiles.

Pedigrees in population biobanks are rarely partitioned into regular families of equal size. Instead, related individuals are typically distributed across small and irregular kinship components. We therefore constructed a second set of cohorts directly from UK Biobank, each containing 10,000 participants, of whom 25%, 50%, or 75% had at least 1 relative within the third degree in the same cohort (Supplementary Method S10). In contrast to the GRAB-generated pedigrees, whitening had little effect in these cohorts.

At a target FDR of 10%, the observed FDRs before whitening were 9.36%, 9.68%, and 9.98% across the three related-participant proportions, compared with 9.44%, 9.46%, and 9.76% after whitening. Power was also nearly unchanged, with the largest absolute difference being only 0.27 percentage points. The limited effect of whitening was consistent with the sparse structure of the GRMs used in these analyses: nonzero off-diagonal entries accounted for only 0.0115%, 0.0127%, and 0.0140% of all off-diagonal entries in the 25%, 50%, and 75% cohorts, respectively. Although many participants had a relative in the cohort, relatedness was confined to small kinship components, leaving the overwhelming majority of pairwise GRM entries equal to zero. Whitened LKOM also retained higher power than SPA_GRM(CCT)_, with power advantages of 1.24%, 2.82%, and 2.58%, respectively (Supplementary Fig. S14).

Together, these results demonstrate that GRM-based whitening enables LKOM to control the FDR under both regular, densely clustered pedigree structures and the sparse, irregular relatedness patterns characteristic of population biobanks, while preserving most of its power and consistently outperforming SPA_GRM(CCT)_.

### LKOM is robust to parameter choices and scales to larger sample sizes

We next examined the sensitivity of LKOM to implementation choices, residual assumptions, and sample size. Relaxing *p*_screen_ from 0.001 to 0.01 increased power by only 0.97%, while increasing runtime per replicate from 195.05 to 340.66 seconds (1.75-fold). A threshold of 0.1 increased power by 2.03% relative to 0.001 but required approximately 4 times the runtime. Tightening the threshold to 0.0001 reduced power by 4.93% while shortening runtime by only 10.4%, from 195.05 to 174.78 seconds (Supplementary Fig. S7). We therefore selected *p*_screen_ = 0.001 to balance power and computational cost. Following KnockoffScreen [18], we pruned variants at *r*^2^ = 0.75 to improve knockoff quality. We assessed sensitivity to this choice by relaxing the threshold to 0.80, 0.85, and 0.90. FDR remained close to the nominal level at 0.80 and 0.85, whereas a threshold of 0.90 increased the observed FDR to 6.8% at *q* = 0.05 and 12.85% at *q* = 0.10 (Supplementary Fig. S9).

Because longitudinal phenotypes may retain outliers and heavy tails after covariate adjustment, we repeated the simulations using variance-standardized *t*_5_ errors. In the mean-only and WS-variability-only settings, observed FDRs were 3.57% and 4.03% at *q* = 0.05, and 9.93% and 9.44% at *q* = 0.10, respectively. In the two-channel setting, the observed FDR also remained controlled, at 4.54% for *q* = 0.05 and 9.95% for *q* = 0.10 (Supplementary Fig. S10).

We further considered shorter longitudinal trajectories, with the number of visits per participant sampled from 2 to 8. The observed FDR of LKOM remained close to the target level, at 4.37% for *q* = 0.05 and 10.97% for *q* = 0.10. Despite the reduced follow-up, LKOM retained the highest channel-wise power among the compared methods at both target levels, reaching 58.72% and 63.14% (Supplementary Tables S2, S3).

LKOM applies a rank-based inverse-normal transformation (INT) to the residual-derived channels to limit the influence of skewness and extreme values. We assessed its contribution by repeating the causal-number and PVE analyses without INT. Omitting INT led to FDR inflation, with a mean observed FDR of 12.92% and a maximum observed FDR of 17.74% at *q* = 0.10. Power averaged across these settings decreased from 68.97% to 54.01% at *q* = 0.05 and from 71.23% to 65.25% at *q* = 0.10 (Supplementary Figs. S11 and S12).

Finally, to assess performance in larger cohorts, we increased the sample size from 10,000 to 100,000 and lowered the minimum MAF from 1.25 *×* 10^*−*3^ to 2 *×* 10^*−*4^. Observed FDR remained below the nominal levels across the 3 nonnull association settings, ranging from 3.19% to 4.81% at *q* = 0.05 and from 7.02% to 8.93% at *q* = 0.10. In the two-channel setting, channel-wise power reached 98.06% and 98.07%, exceeding TrajGWAS by 6.11% and 5.19%, respectively (Supplementary Fig. S13). These results support the applicability of LKOM to larger cohorts containing rarer variants. We further evaluated the computational scalability of LKOM across sample sizes from 10,000 to 100,000. All benchmarks were conducted using the same computing environment with eight CPU threads. Mean runtime per simulation replicate increased from 5.06 to 23.40 min, compared with 1.06 to 5.75 min for the TrajGWAS-based analysis. Marginal screening accounted for the largest proportion of the LKOM runtime at larger sample sizes, while peak memory increased from 6.25 to 34.55 GB (Supplementary Fig. S15). Both runtime and peak memory showed empirical scaling close to *O*(*n*) across the evaluated sample sizes, suggesting that the computational demands of LKOM remain manageable as *n* increases.

## Discussion

In this study, we developed LKOM, a knockoff-based framework for jointly identifying genetic effects on the mean and WS variability of longitudinal phenotypes. LKOM combines subject-level responses derived from a longitudinal location–scale model with a dual-channel Bayesian working model and masked likelihood-ratio statistics. This construction provides variant-level FDR control for the joint null of no association with either channel, while further characterizing selected variants as mean-specific, WS-variability-specific, or shared. Across simulations based on UK Biobank genotypes, LKOM maintained observed FDR close to the target levels under diverse genetic architectures and generally achieved greater power than separate-channel knockoff and marginal association approaches. The largest gains occurred for shared effects and rare variants, particularly when the causal variants collectively explained a smaller proportion of phenotypic variation. A GRM-based whitening extension further enabled LKOM to accommodate related samples.

A central feature of LKOM is that it models the mean and WS variability channels jointly without requiring their genetic architectures to be identical. Previous longitudinal-trait analyses have shown that loci affecting WS variability frequently also influence mean levels [2, 7]. The four-state model used by LKOM allows such shared effects while retaining separate mean-only and variability-only states. Consistent with this motivation, the advantage of LKOM over LKOM-single increased markedly with the proportion of causal variants shared between the two channels and was greatest under low PVE, where the evidence available from either channel alone was limited. By contrast, the detection of mean-only variants was largely preserved, suggesting that borrowing information across channels did not require a substantial sacrifice in sensitivity to channel-specific effects. The state-classification results provided a similar picture that most detected variants were assigned to their true states, whereas shared variants were occasionally classified as mean-only under weak signals. This asymmetry is consistent with the greater difficulty of detecting effects on WS variability and diminished as signal strength increased. These findings indicate that the gain from joint modeling arises primarily when complementary evidence from the two channels can reinforce an otherwise weak association.

The inferential target of LKOM differs from that of conventional longitudinal GWAS. Marginal association tests evaluate each variant separately and may therefore identify variants that tag an associated variant through linkage disequilibrium. LKOM instead evaluates whether a variant contributes information conditional on the other variants included in the analysis. This multivariable target can improve localization within correlated regions, although a conditionally selected variant should not automatically be interpreted as causal. LKOM also differs from simply applying knockoff selection separately to the two channels. Separate analyses do not exploit shared association patterns, and combining their discoveries by union does not in general provide the same FDR guarantee. In LKOM, the masked likelihood-ratio working model is used to distinguish the original member of each original–knockoff pair by integrating evidence across both channels [19]. Under valid model-X knockoffs, FDR control follows from pairwise exchangeability, conditional independence and the flip-sign property of the resulting statistic [15]. The Bayesian working model therefore primarily determines statistical efficiency and state classification rather than the validity of the knockoff filter. The posterior state assignments should accordingly be regarded as post-selection characterizations, and they do not carry a separate state-specific FDR guarantee.

The comparison methods considered here also operate under different selection criteria. LKOM targets variant-level FDR for the dual-channel joint null, whereas LKOM-single and TrajKnock apply knockoff filtering separately within each channel, and TrajGWAS and SPA_GRM_ use channel-specific multiplicity correction. These differences are partly unavoidable because the comparison methods were designed for marginal or single-channel analyses. The power comparisons should therefore be interpreted as comparisons of discovery performance under each method’s intended selection rule rather than as evidence of universal dominance under an identical error criterion. Nevertheless, the comparison with LKOM-single directly isolates the contribution of joint modeling because it uses the same underlying response construction and MLR framework. The increasing advantage of LKOM over LKOM-single with greater cross-channel sharing provides evidence that the observed power gains are attributable to the dual-channel model rather than to knockoff selection alone.

LKOM was robust across several departures from the primary simulation design. Its FDR remained close to the target levels under variance-standardized heavy-tailed errors and when participants had only 2 to 8 longitudinal measurements. The latter result is relevant to electronic health records, in which follow-up intensity commonly differs across participants and many individuals have relatively few observations. The rank-based inverse-normal transformation was an important component of this robustness. Omitting it produced appreciable FDR inflation and reduced power, particularly through instability in the WS variability channel. Because the transformation changes the scale of the residual-derived responses, however, LKOM is primarily a discovery and classification procedure rather than a method for estimating genetic effects on the original phenotype scale. The symmetric marginal screen and LD pruning also represent important practical choices. Marginal screening substantially reduced computation while preserving the swap symmetry required by the knockoff statistic, whereas overly permissive LD pruning weakened empirical FDR control.

The effect of sample relatedness depended strongly on the structure of the GRM. In simulations with regular, densely connected pedigrees, ignoring between-subject dependence inflated FDR, whereas GRM-based whitening restored control with only a modest reduction in power. In cohorts constructed directly from UK Biobank, whitening had little effect even when a large proportion of participants had a relative in the cohort. These cohorts contained many small and sparsely connected kinship components, so the proportion of related participants did not translate into dense covariance across the sample. This contrast indicates that the density and topology of relatedness can be more informative than the percentage of participants classified as related when assessing its likely effect on longitudinal association analyses. Whitened LKOM also retained higher power than SPA_GRM(CCT)_ in the simulated settings. The two approaches nevertheless address different objectives: SPA_GRM_ provides calibrated marginal association tests across genotype and phenotype distributions [7], whereas LKOM performs multivariable dual-channel selection with FDR control. They may therefore serve complementary roles in longitudinal genetic analysis.

Several limitations should be considered. First, the theoretical guarantee of LKOM depends on the validity of the model-X knockoffs. Misspecification of the genotype distribution or insufficient exchangeability between original and knockoff genotypes may compromise FDR control, particularly for rare variants or in regions with complex LD. Second, the current implementation uses LD pruning and marginal screening to make posterior computation feasible. Although the screening rule is symmetric, weak variants that do not pass the screen cannot subsequently be recovered, and aggressive pruning may reduce genomic resolution. Third, the current framework summarizes longitudinal phenotypes through mean and WS variability channels. Other trajectory features, including longitudinal slopes, nonlinear change and responses to clinical interventions, are not explicitly modeled. Fourth, the simulations used UK Biobank-derived genotypes and relatedness structures, but performance may differ across ancestries, genotyping platforms and patterns of longitudinal follow-up. Real-data applications are required to establish the biological interpretability and reproducibility of the additional discoveries. Finally, posterior computation remains more demanding than marginal testing. Further improvements in blockwise computation, parallelization and software implementation will be important for routine genome-wide use.

In summary, LKOM integrates model-X knockoffs, masked likelihood-ratio statistics and joint modeling of longitudinal mean and WS variability into a unified variable-selection framework. Its principal advantage arises when genetic effects are shared across the two channels, especially when individual signals are weak or carried by rarer variants. By providing FDR-controlled conditional discoveries together with interpretable association-state summaries, LKOM offers a framework for extracting genetic information from longitudinal phenotypes that is not captured by analyses of mean levels alone.

## Methods

### Longitudinal location–scale mixed-effects model

We adopt the longitudinal location–scale mixed-effects model [2] to characterize genetic effects on both the mean and within-subject (WS) variability of a longitudinal trait. For subject *i* = 1, …, *m* at measurement occasion *j* = 1, …, *n*_*i*_, the mean submodel is

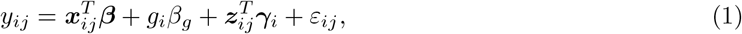

where *y*_*ij*_ is the observed trait value, 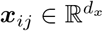 is a vector of fixed-effect covariates with coefficients ***β***, and 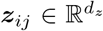 is a vector of random-effect covariates with subject-specific coefficients ***γ***_*i*_. For exposition, *g*_*i*_ denotes the allele dosage of a candidate variant for subject *i*, and *β*_*g*_ represents its effect on the trajectory mean. Conditional on the random effects, the residual term satisfies E(*ε*_*ij*_) = 0, with visit-specific variance

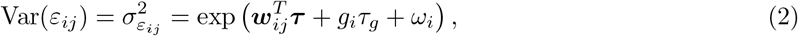

where 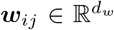 contains covariates for the variance submodel, ***τ*** denotes their fixed effects, *τ*_*g*_ is the genetic effect on WS variability, and *ω*_*i*_ is a subject-specific random intercept in the variance submodel. Thus, *β*_*g*_ characterizes a shift in the mean trajectory, whereas *τ*_*g*_ characterizes a multiplicative change in the residual variance on the log scale. The mean- and variance-level random effects are assumed to have mean zero and covariance structure

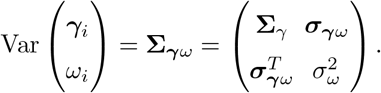

The random effects are independent across subjects and are assumed to be independent of the residual errors. The number and timing of measurements may vary across subjects. The covariate vectors may contain both time-invariant variables, such as sex and genetic ancestry principal components, and time-varying variables, such as age and medication status. Intercept terms may also be included as appropriate in the fixed- and random-effect components.

The two genetic-effect parameters define four possible association states for a variant: no effect (*β*_*g*_ = 0, *τ*_*g*_ = 0), a mean effect only (*β*_*g*_*≠* 0, *τ*_*g*_ = 0), a WS variability effect only (*β*_*g*_ = 0, *τ*_*g*_*≠* 0), or effects on both components (*β*_*g*_*≠* 0, *τ*_*g*_*≠* 0). Accordingly, LKOM performs variant selection with respect to the joint null hypothesis

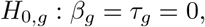

such that a selected variant may affect the trajectory mean, WS variability, or both. Although *g*_*i*_ denotes a single variant in Equations (1) and (2), LKOM subsequently models multiple original and knockoff variants jointly and uses the posterior probabilities of the four states to characterize their association patterns.

### WiSER-based construction of subject-level responses

LKOM fits the null longitudinal location–scale model once using WiSER [6], excluding genetic effects from both submodels by imposing *H*_0_ : *β*_*g*_ = *τ*_*g*_ = 0. The fitted null model is shared across all subsequent variant analyses. Using this fit, LKOM constructs subject-level score contributions for the mean and WS variability channels. Let

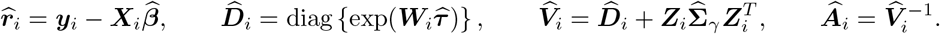

The two score contributions are

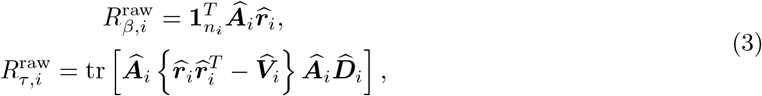

where 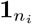 is an *n*_*i*_-dimensional vector of ones. These quantities are proportional to the individual contributions to the WiSER score statistics for testing genetic effects on the mean and WS variability, respectively.

To reduce the influence of extreme score contributions and improve the numerical stability of the downstream working model, a rank-based inverse normal transformation is applied separately to the two channels. For *k ∈ {β, τ }*,

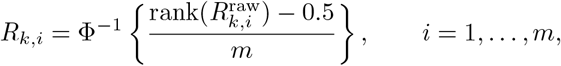

where Φ^*−*1^ is the standard normal quantile function and average ranks are used for ties. The transformed vectors ***R***_*β*_ and ***R***_*τ*_ form the two response channels in the subsequent masked likelihood ratio model.

### Knockoff construction

The model-X knockoff framework [15] controls the false discovery rate (FDR) in variable selection by constructing synthetic negative-control variables. Given the genotype matrix ***G*** *∈* ℝ^*m×p*^, a knockoff copy 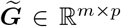 is generated without using the phenotype. A valid knockoff construction satisfies pairwise exchangeability,

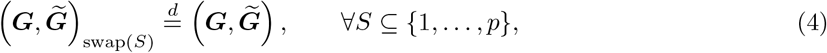

together with conditional independence between the knockoffs and the response given the original genotypes. Here, swap(*S*) exchanges *G*_*j*_ and 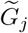 for every *j ∈ S*. Consequently, the knockoff variables preserve the dependence structure among genetic variants while serving as negative controls for assessing conditional associations.

To generate knockoff genotypes efficiently, LKOM adopts the sequential conditional independent pairs (SCIP) strategy implemented in KnockoffScreen [18]. At step *j*, the knockoff of variant *j* is sampled according to

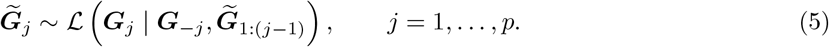

To reduce the computational burden, this conditional distribution is approximated using variants in a local linkage disequilibrium neighborhood *B*_*j*_:

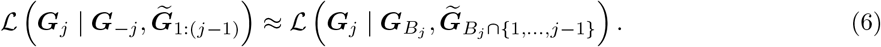

The corresponding conditional mean is modeled as

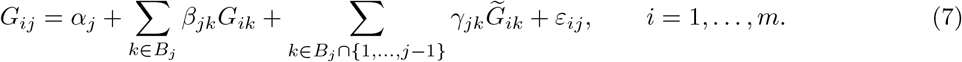

After estimating the regression parameters, let *Ĝ*_*ij*_ denote the fitted value and 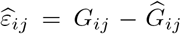 the corresponding residual. The knockoff genotype is constructed by randomly permuting the residuals across subjects:

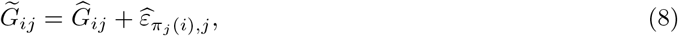

where *π*_*j*_ denotes a random permutation of *{*1, …, *m}*. The resulting augmented genotype matrix [***G***, 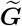] is subsequently incorporated into the masked likelihood ratio model to construct knockoff statistics for variant selection.

### Bayesian dual-channel masked likelihood ratio model

Following the masked likelihood ratio (MLR) framework of Spector and Fithian [19], LKOM jointly models the two WiSER-derived response channels to distinguish each original variant from its knockoff while accounting for uncertainty in its association pattern. Let ***Y*** = [***R***_*β*_, ***R***_*τ*_ ] *∈* ℝ^*m×*2^ denote the response matrix. Before model fitting, the response and genotype columns are centered and scaled to unit empirical variance.

For each original–knockoff pair, a deterministic ordering rule (Supplementary Method S1) based only on the two genotype vectors produces a canonical pair (***a***_*j*_, ***b***_*j*_). This ordering does not depend on the phenotype or the original–knockoff labels. Bayesian inference uses the masked data

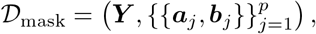

with the observed original–knockoff identities withheld. A latent orientation variable *S*_*j*_ *∈ {A, B}* specifies which member of pair *j* enters the working regression:

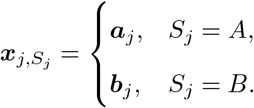

Here, *p* denotes the number of variant pairs included in the joint fit.

The dual-channel working model is

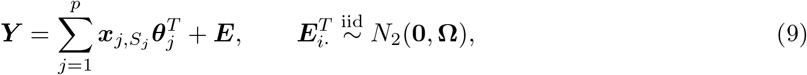

where ***θ***_*j*_ = (*θ*_*βj*_, *θ*_*τj*_)^*T*^ contains the regression coefficients for the mean and WS variability response channels, respectively. These coefficients are defined on the transformed response scale and are distinct from the genetic-effect parameters in the longitudinal model. The residual covariance matrix **Ω** accommodates dependence between the two response channels.

To represent channel-specific and shared associations, we introduce *H*_*j*_ *∈ ℋ* = *{*00, 10, 01, 11*}*, with active coordinate sets *A*_00_ = ∅, *A*_10_ = *{*1*}, A*_01_ = *{*2*}*, and *A*_11_ = *{*1, 2*}*. We assign equal prior probabilities to the two orientations and a hierarchical prior to the association states:

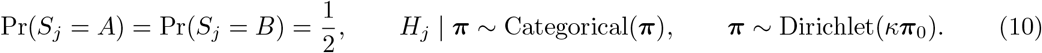

Conditional on *H*_*j*_ = *h*, inactive coefficients are fixed at zero and active coefficients follow a Gaussian slab:

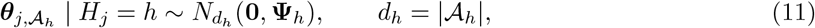

where ***θ***_*j*_ = **0** for *h* = 00, and

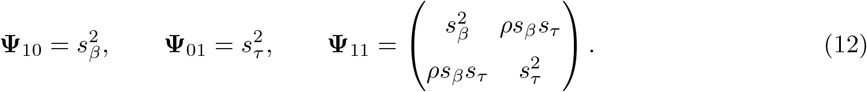

The parameters *s*_*β*_ and *s*_*τ*_ determine the channel-specific slab scales, whereas *ρ* describes the prior correlation of effects in the shared state.

We place an inverse-Wishart prior **Ω** *∼* IW_2_(*ν*_0_, ***S***_0_). The slab scales and correlation are assigned discrete priors on prespecified grids *G*_*β*_, *G*_*τ*_ *⊂* (0, *∞*) and *G*_*ρ*_ *⊂* (*−*1, 1). Prior masses for *s*_*k*_, *k ∈ {β, τ }*, are proportional to a half-*t* density evaluated on *G*_*k*_, and those for *ρ* are proportional to 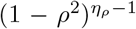, corresponding to a discretized two-dimensional LKJ correlation prior with shape parameter *η*_*ρ*_. Posterior inference is performed using a Gibbs sampler. Conditional on the current effect coefficients and association states, *s*_*β*_, *s*_*τ*_, and *ρ* are updated sequentially by enumerating their discrete full conditional distributions over the prespecified grids and sampling from the resulting normalized weights.

For variant pair *j*, define the partial residual

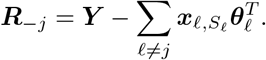

Conditional on the remaining parameters, the active coefficients can be integrated out analytically to obtain a conditional Bayes factor for each orientation–state combination:

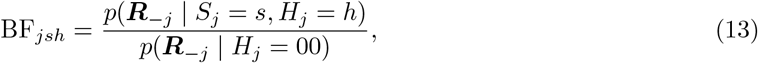

with BF_*js*,00_ = 1. Specifically, writing

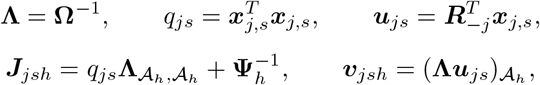

the conditional Bayes factor for *h≠* 00 is

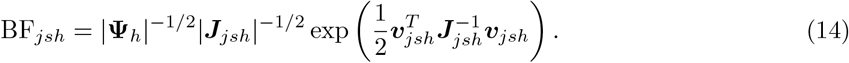

The eight orientation–state combinations are sampled with normalized probabilities

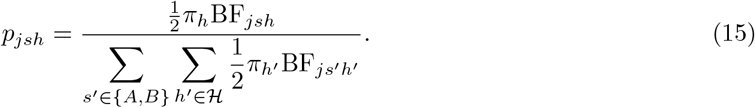

After sampling (*S*_*j*_, *H*_*j*_) = (*s, h*), the active effects are drawn from

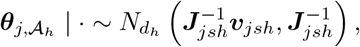

while inactive effects remain zero.

After each sweep over the variant pairs, ***π*** and **Ω** are updated from their Dirichlet and inverse-Wishart full conditional distributions. The sampler can be initialized with all effects set to zero, and a deterministic conditional-Bayes-factor fit is available as an optional warm start. The complete likelihood and posterior distribution, the derivation of the conditional Bayes factor, the hyperparameter settings, and the optional initialization procedure are described in the Supplementary Method S2-S5.

Following burn-in, posterior orientation and state probabilities are estimated by Rao–Blackwellized averages over *M* retained iterations:

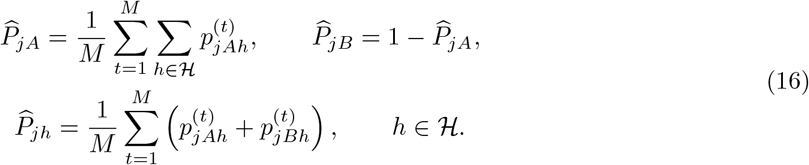

The orientation probabilities quantify the evidence distinguishing the two members of each masked pair and provide the posterior odds used to construct the MLR statistic. The state probabilities summarize support for the four channel-association patterns after marginalizing over orientation.

### MLR statistics and variable selection

Before joint Bayesian model fitting, LKOM applies pairwise-symmetric marginal screening. For independent samples, marginal association *p*-values are obtained using TrajGWAS, whereas SPA_GRM_ is used for related samples to account for between-subject dependence. When computational speed is prioritized, either procedure can be replaced by the simpler Norm test, which uses the conventional *t*-test *p*-value for the genotype coefficient in a marginal linear regression. In all cases, the same screening procedure is applied separately to the original and knockoff genotypes in both the mean and WS variability channels. Let *p*_*βj*_, *p*_*τj*_ denote the two *p*-values for the original genotype and 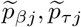 their knockoff counterparts. The set of variant pairs entering the dual-channel MLR model is

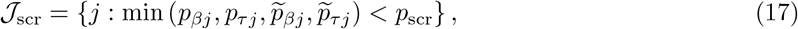

where *p*_scr_ is a prespecified screening threshold. The entire original–knockoff pair is retained whenever either member shows marginal evidence of association in either response channel. This rule is unchanged by exchanging the original and knockoff variables and therefore preserves pairwise symmetry. Variants not retained by the screening step are assigned *W*_*j*_ = 0.

For each retained pair, let 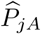 and 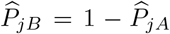 denote the posterior probabilities that the active member of the canonical pair is ***a***_*j*_ or ***b***_*j*_, respectively. The canonical posterior log-odds is

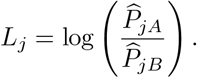

Let *o*_*j*_ = 1 if the observed original genotype corresponds to ***a***_*j*_, and *o*_*j*_ = *−*1 if it corresponds to ***b***_*j*_. The LKOM feature statistic is defined as

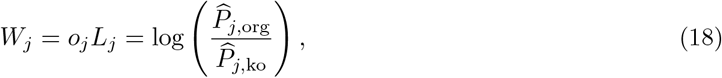

where 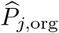 and 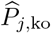 are the posterior probabilities assigned to the original and knockoff members of pair *j*. Accordingly, *W*_*j*_ *>* 0 indicates stronger evidence for the original variant, whereas *W*_*j*_ *<* 0 indicates stronger evidence for its knockoff; the magnitude |*W*_*j*_| reflects the posterior confidence in distinguishing the two members of the masked pair.

Because the canonical ordering is invariant to exchanging the original and knockoff labels, such an exchange reverses *o*_*j*_ while leaving *L*_*j*_ unchanged. Consequently,

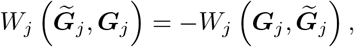

which establishes the antisymmetry required of a valid knockoff statistic. A proof of the invariance of the masked representation under original–knockoff swaps, together with the resulting antisymmetry of the MLR statistic, is provided in the Supplementary Method S6.

At a target FDR level *q*, define the set of nonzero candidate thresholds as

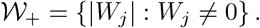

The knockoff+ threshold [15] is

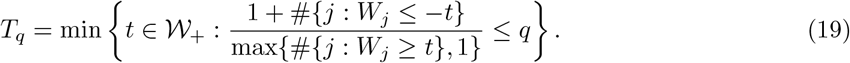

If no candidate threshold satisfies this condition, no variant is selected. Otherwise, LKOM reports

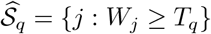

as the set of discoveries against the joint null hypothesis *H*_0,*j*_ : *β*_*j*_ = *τ*_*j*_ = 0.

For each selected variant, LKOM additionally reports the posterior probabilities 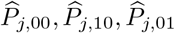, and 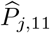, obtained by marginalizing over the latent orientation. These probabilities characterize the evidence for no association, a mean-only association, a WS-variability-only association, and an association with both channels, respectively. A maximum-posterior summary may be defined as

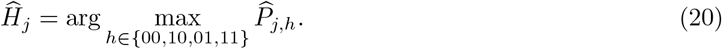

This state characterization is used to interpret the association pattern of a selected variant and does not enter the knockoff+ thresholding procedure.

### Related samples

The model above assumes independence across subjects. In samples containing related individuals, genetic sharing induces between-subject dependence that, if ignored, may inflate type I error rates [7]. Let **Φ** *∈* ℝ^*m×m*^ denote the standardized genetic relationship matrix (GRM), aligned with the *m* study subjects. A latent-effect representation of this dependence augments the longitudinal location–scale model as

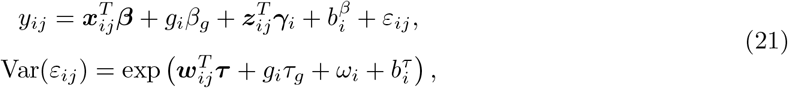

where

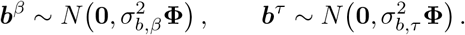

LKOM represents the resulting between-subject dependence through a GRM-structured row covariance for the bivariate response matrix. Rather than refitting the expanded longitudinal model for each variant, LKOM whitens the WiSER-derived response matrix and the augmented original–knockoff genotype matrix before fitting the dual-channel MLR model.

Let

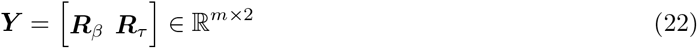

denote the bivariate response matrix. We assume the working model

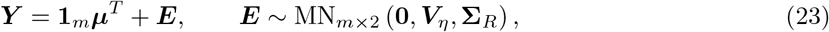

where **Σ**_*R*_ *∈* ℝ^2*×*2^ is the working covariance between the mean and WS variability responses in the relatedness model. The covariance across subjects is parameterized as

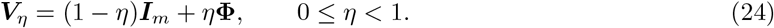

Thus, *η* = 0 corresponds to independent subjects, whereas larger values of *η* assign greater weight to the dependence structure represented by the GRM.

Conditional on *η*, generalized least-squares centering and whitening are applied jointly to the response and augmented genotype matrices. For

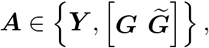

the transformed matrix is

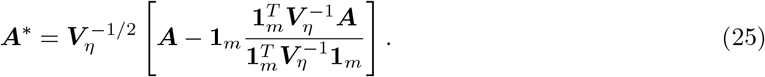

The fraction in Equation (25) is the vector of generalized least-squares column means. The transformation removes the intercept and adjusts for between-subject dependence under the fitted GRM-based covariance model. The remaining covariance between the two response channels is accommodated by the dual-channel MLR likelihood. With the fitted covariance held fixed, applying the same transformation to the original and knockoff genotype matrices preserves symmetry under within-pair column swaps.

LKOM estimates *η* once from the bivariate response matrix under a null model containing no genetic effects. For a candidate value of *η*, let 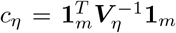. The generalized least-squares estimator of the two-channel intercept vector is

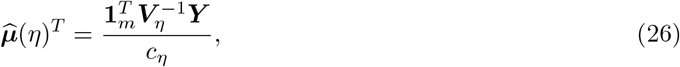

with residual matrix

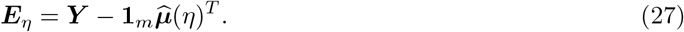

The corresponding restricted maximum-likelihood estimator of the between-channel covariance is

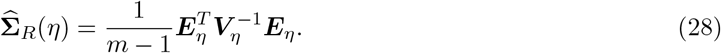

After profiling out both ***µ*** and **Σ**_*R*_, the profile restricted negative log-likelihood for *η*, up to an additive constant, is

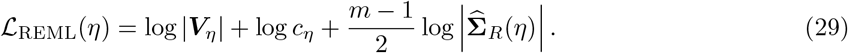

The relatedness parameter is estimated by the one-dimensional optimization

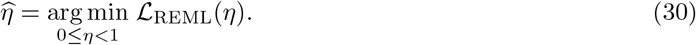

Using 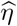, LKOM sets 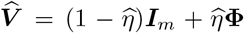 and applies the transformation in Equation (25), with *η* replaced by 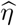, to the response, original-genotype, and knockoff-genotype matrices. Because 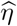 is estimated only from ***Y*** and **Φ**, and the same transformation is applied to both members of each pair, exchanging an original variant with its knockoff commutes with whitening. Applying the same whitening transformation to both members of each pair preserves the swap symmetry of the feature-statistic construction. The profile REML derivation are provided in the Supplementary Method S7.

### Simulation studies

#### Genotype and knockoff data

We used real genotypes from UK Biobank to preserve the allele-frequency spectrum and local linkage disequilibrium (LD) structure. For the primary simulations, 10,000 unrelated White British participants were sampled using a fixed random seed 718. Starting from the quality-controlled imputed genotype data, variants were required to have genotype missingness no greater than 2%, Hardy–Weinberg equilibrium test *p ≥* 10^*−*6^, and minor allele count at least 25 (minor allele frequency (MAF) at least 1.25 *×* 10^*−*3^). Variants were LD pruned at *r*^2^ = 0.75. The genotype was partitioned into LD blocks, and five consecutive blocks were randomly selected in each replicate. One knockoff copy was generated for every retained variant using `create.MK` in the R package `KnockoffScreen` (version 0.3.0). Original and knockoff genotypes were kept in matched pairs throughout screening and model fitting.

#### Longitudinal phenotype generation

Longitudinal outcomes were generated from the location–scale model in Equations (1) and (2). Unless otherwise specified, the number of measurements for each subject was sampled uniformly from *{*6, …, 15*}*. The mean and variance covariate matrices were ***X*** = ***W*** = (1, *x*_1_, *x*_2_, *x*_3_), ***Z*** = (1, *z*), where *x*_1_ *∼* Bernoulli(0.5) and *x*_2_ *∼ N* (0, 0.5^2^) were generated at the subject level, whereas *x*_3_ *∼ N* (0, 0.3^2^) and *z ∼ N* (0, 0.5^2^) were generated at the measurement level. We set

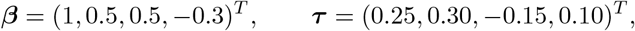

and generated (*γ*_0*i*_, *γ*_1*i*_, *ω*_*i*_)^*T*^ independently from a mean-zero multivariate normal distribution with covariance matrix 0.5***I***_3_. Conditional residuals were normally distributed in the primary simulations, with variance determined by the variance submodel.

Let *C*_*β*_ and *C*_*τ*_ denote the sets of variants affecting the mean and WS variability channels. Causal variants were sampled from the variants in the selected blocks. For channel *c ∈ {β, τ }* and causal variant *k ∈ C*_*c*_, the initial effect magnitude was

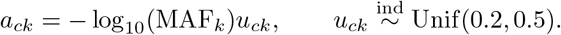

The corresponding unscaled subject-level genetic score was

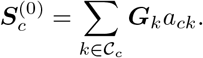

The resulting subject-level genetic scores were rescaled to attain prespecified channel-specific PVE. The scale factors were

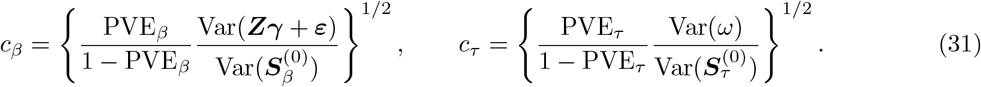

For the mean channel, the genetic score was repeated over the observed measurements before calculating its empirical variance. The final genetic effects were *c*_*β*_*a*_*βk*_ and *c*_*τ*_ *a*_*τk*_, respectively. Thus, the target PVE was held fixed when the number or MAF distribution of causal variants changed.

#### Primary simulation scenarios

Unless otherwise specified, simulations included 20 causal variants in each channel, with (PVE_*β*_, PVE_*τ*_ ) = (0.05, 0.10). The two causal sets were sampled independently, with no imposed overlap. The default LD-pruning and marginal screening thresholds were *r*^2^ = 0.75 and *p*_scr_ = 0.001, respectively.

We first examined performance across the four association states defined by LKOM. The null state contained no causal variants, the mean-only and WS-variability-only states contained 20 causal variants in the active channel, and the both channels state contained 20 variants in each channel. For each nonnull state, we considered (PVE_*β*_, PVE_*τ*_ ) *∈ {*(0.025, 0.05), (0.05, 0.10), (0.10, 0.20)*}*, with the PVE of an inactive channel set to zero.

To assess the effects of signal sparsity and cross-channel sharing, we varied the number and overlap of causal variants. The causal-number analysis used |*C*_*β*_| = |*C*_*τ*_ | *∈ {*5, 20, 50*}*, crossed with the three PVE pairs above. We defined the overlap proportion as *ω* = (|*C*_*β*_ *∩ C*_*τ*_ |)*/* min(|*C*_*β*_|, |*C*_*τ*_ |) . The overlap analysis used 20 causal variants per channel and varied *ω* over 0, 0.2, 0.5, 0.8, and 1, crossed with the three PVE pairs. When *ω* = 0, the two causal sets were sampled independently with no imposed sharing, so incidental overlap could occur by chance.

Robustness to asymmetric genetic architectures was examined by fixing one channel at 20 causal variants and varying the other over 5, 10, and 15, in both directions. We also considered the unequal PVE combinations (0.025, 0.10), (0.025, 0.20), (0.05, 0.05), (0.05, 0.20), (0.10, 0.05), (0.10, 0.10), with 20 causal variants in each channel.

#### Related-sample simulations

Two related-sample designs were used to evaluate the effect of GRM-based whitening under different forms of family structure. In the first, family-structured genotypes were generated from UK Biobank data using `GRAB.SimuGMatFromGenoFile` in the R package `GRAB` (version 0.2.4). The three configurations combined 5,000 unrelated individuals with 1,250 four-member families, 500 ten-member families, or 250 twenty-member families, giving 10,000 individuals in each setting. In the second design, 10,000 UK Biobank participants were sampled so that 25%, 50%, or 75% had at least one third-degree-or-closer relative in the analysis cohort. Genotype blocks, knockoffs, and the GRM were constructed separately within each cohort.

For both designs, relatedness-induced effects were generated as

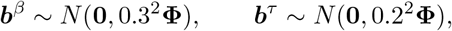

and added to the mean and log-variance submodels, respectively. All other settings followed the default specification. LKOM was applied with and without GRM-based whitening. To isolate the effect of whitening, the variant pairs retained by the unwhitened marginal screening step were reused in the whitened analysis.

#### Additional sensitivity analyses

Sensitivity analyses examined the effects of preprocessing, posterior computation, tuning parameters, distributional assumptions, follow-up length, and sample size. Each analysis varied one component of the default setting unless otherwise specified. The contribution of the inverse normal transformation was evaluated by repeating the balanced causal-number–PVE scenarios without transforming the two WiSER-derived response channels. We also replaced posterior sampling with the deterministic conditional-Bayes-factor procedure and varied

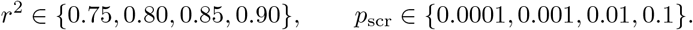

Robustness to non-Gaussian residuals was assessed using variance-standardized *t*_5_ innovations. Reduced follow-up was examined by sampling *n*_*i*_ uniformly from *{*2, …, 8*}* rather than *{*6, …, 15*}*. Finally, scalability and performance for rare and low-frequency variants were evaluated in 100,000 participants with a minimum MAF of 2 *×* 10^*−*4^. Full scenario combinations and computational settings are provided in the Methods and Supplementary Table S1.

#### Evaluation

The primary comparisons included dual-channel LKOM, LKOM-single, TrajGWAS, and TrajKnock. LKOM-single fitted the single-channel MLR model separately to the mean and WS variability responses and applied the knockoff+ filter at level *q* within each channel. Let *Ŝ*_*β*_ and *Ŝ*_*τ*_ denote the resulting channel-specific discovery sets. The overall LKOM-single discovery set was defined as *Ŝ*_*β*_ *∪ Ŝ*_*τ*_ .

TrajGWAS evaluated marginal associations separately for the mean and WS variability channels. Let *p*_*g*_ denote the total number of SNP–replicate hypotheses within each channel in grouped evaluation *g*, as described below. A hypothesis was declared significant in channel *k ∈ {β, τ }* when its marginal *p*-value satisfied

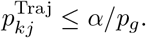

For each comparison, we set *α* = *q*. The two channel-specific discovery sets were combined by union. TrajKnock applied TrajGWAS separately to the original and knockoff genotypes and constructed the channel-specific statistic

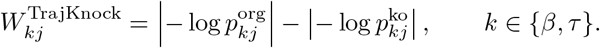

The knockoff+ filter was applied at level *q* separately within each channel, after which the two discovery sets were combined by union. In the related-sample simulations, we additionally used the best-performing SPA_GRM_+CCT method reported by Xu et al. [7], combining the *p*-values from the untransformed and inverse-normal-transformed responses within each channel, thresholding the resulting *p*-values at *α/p*_*g*_, and combining the two channel-specific discovery sets by union.

No additional multiplicity adjustment was applied when combining the channel-specific discoveries from the comparison methods. LKOM targeted variant-level FDR for the joint null, whereas LKOM-single and TrajKnock applied knockoff+ separately within each channel, and TrajGWAS and SPA_GRM_+CCT used channel-specific Bonferroni correction. Power was compared under these method-specific selection criteria.

For each scenario, we generated 200 independent simulation replicates and constructed 500 grouped evaluations. Each group contained 5 distinct replicates sampled without replacement from the 200 datasets; individual replicates could be reused across groups. Models were fitted separately within each replicate. Within each group, SNP-level statistics were pooled across the 5 replicates, and knockoff+ thresholds were recalculated jointly for LKOM and separately within each channel for LKOM-single and TrajKnock. This procedure reduced the impact of the minimum discovery count required by knockoff+, which is at least *⌈*1*/q⌉*, even when no negative statistics are observed. For TrajGWAS and SPA_GRM_+CCT, Bonferroni correction used *p*_*g*_ hypotheses within each channel. FDR and power were calculated for each group and averaged across the 500 evaluations. These evaluations were resamples of the 200 independently generated datasets, rather than additional independent simulations.

Let *ℬ*_*g*_ denote the five replicates included in grouped evaluation *g*, and let *Ŝ*_*gr*_ be the overall discovery set for replicate *r ∈ ℬ*_*g*_ under the corresponding grouped threshold. The false discovery proportion was calculated against the joint null as

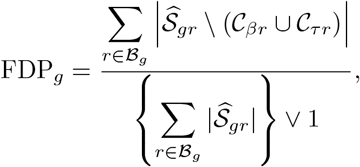

where *C*_*βr*_ and *C*_*τr*_ denote the true causal sets in replicate *r*. Empirical FDR was estimated by

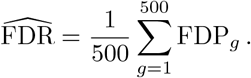

This measure evaluates false discoveries against the joint null and does not quantify errors in channel attribution or association-state classification.

Channel-wise power was defined as the equal-weight average of the two channel-specific detection rates. For grouped evaluation *g*,

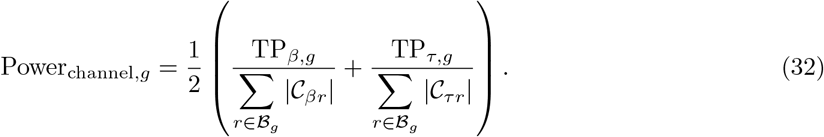

In scenarios with causal variants in only one channel, power was calculated for that channel; power was not defined under the global null.

For dual-channel LKOM, the channel-specific true-positive counts were

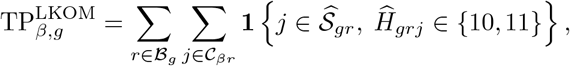

and

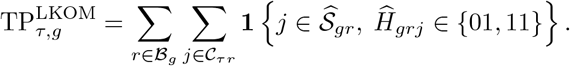

Thus, a causal mean-effect variant was counted as a true positive for the mean channel only when it was selected and assigned to state 10 or 11, whereas a causal WS-variability-effect variant was counted for the variability channel only when it was selected and assigned to state 01 or 11.

For LKOM-single, TrajGWAS, TrajKnock, and SPA_GRM_+CCT, the corresponding counts were

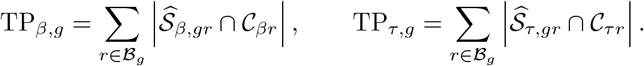

A variant affecting both channels was included in both channel-specific denominators and contributed to each numerator only when detected in the corresponding channel. Reported channel-wise power was the average of Power_channel,*g*_ over the 500 grouped evaluations.

Empirical FDR was evaluated over target levels *q* = 0.01, …, 0.10, with power primarily reported at *q* = 0.05. Power was also summarized within rare (MAF *<* 0.01), low-frequency (0.01 *≤* MAF *<* 0.05), and common (MAF *≥* 0.05) causal-variant strata by restricting the true-positive counts and causal-set denominators to the corresponding MAF category.

To assess the recovery of complete association states, we additionally evaluated exact-state power. For methods with separate channel-specific discovery sets, the predicted state was defined by membership in those sets: 10 for mean-only detection, 01 for WS-variability-only detection, 11 for detection in both channels, and 00 for detection in neither. Let *H*_*rj*_ denote the true association state of variant *j* in replicate *r*. Exact-state power was defined as

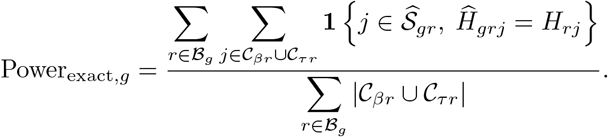

Each causal variant was counted once, including variants affecting both channels. Reported exact-state power was averaged over the 500 grouped evaluations.

To assess classification conditional on detection, we constructed a count 4 *×* 4 confusion matrix. Suppressing the group and replicate indices and letting *j* index variant–replicate pairs, the corresponding entry was

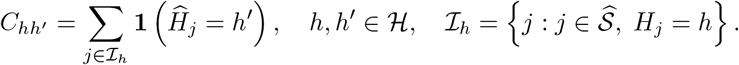

Each row gives the proportions of detected variants from a given true state assigned to the four possible states, with diagonal entries representing correct post-selection classification. Unlike exact-state power, this matrix excludes undetected causal variants.

## Supplementary Material

Supplementary Material is available for this paper and includes Supplementary Methods S1–S10, Supplementary Figs. S1–S15, and Supplementary Tables S1–S9. Supplementary Tables S2–S4, S6, S8, and S9 are provided as a separate Excel file.

## Code availability

The LKOM R package, source code, documentation, and installation instructions are publicly available at https://github.com/hyd020718/LKOM.

## References

[1] Bycroft, C., Freeman, C., Petkova, D., et al.: The UK Biobank resource with deep phenotyping and genomic data. Nature 562(7726), 203–209 (2018) 10.1038/s41586-018-0579-z

[2] Ko, S., German, C.A., Jensen, A., et al.: GWAS of longitudinal trajectories at biobank scale. The American Journal of Human Genetics 109(3), 433–445 (2022) 10.1016/j.ajhg.2022.01.018

[3] Gorst, C., Kwok, C.S., Aslam, S., et al.: Long-term glycemic variability and risk of adverse outcomes: A systematic review and meta-analysis. Diabetes Care 38(12), 2354–2369 (2015) 10.2337/dc15-1188

[4] Rothwell, P.M., Howard, S.C., Dolan, E., et al.: Prognostic significance of visit-to-visit variability, maximum systolic blood pressure, and episodic hypertension. The Lancet 375(9718), 895–905 (2010) 10.1016/S0140-6736(10)60308-X

[5] Almuwaqqat, Z., Hui, Q., Liu, C., et al.: Long-term body mass index variability and adverse cardiovascular outcomes. JAMA Network Open 7(3), 243062 (2024) 10.1001/jamanetworkopen.2024.3062

[6] German, C.A., Sinsheimer, J.S., Zhou, J., Zhou, H.: WiSER: Robust and scalable estimation and inference of within-subject variances from intensive longitudinal data. Biometrics 78(4), 1313–1327 (2022) 10.1111/biom.13506

[7] Xu, H., Ma, Y., Xu, L.-l., et al.: SPA_GRM: Effectively controlling for sample relatedness in large-scale genome-wide association studies of longitudinal traits. Nature Communications 16, 1413 (2025) 10.1038/s41467-025-56669-1

[8] Schaid, D.J., Chen, W., Larson, N.B.: From genome-wide associations to candidate causal variants by statistical fine-mapping. Nature Reviews Genetics 19(8), 491–504 (2018)

[9] Uffelmann, E., Huang, Q.Q., Munung, N.S., De Vries, J., Okada, Y., Martin, A.R., Martin, H.C., Lappalainen, T., Posthuma, D.: Genome-wide association studies. Nature reviews methods primers 1(1), 59 (2021)

[10] Watanabe, K., Taskesen, E., Van Bochoven, A., Posthuma, D.: Functional mapping and annotation of genetic associations with fuma. Nature communications 8(1), 1826 (2017)

[11] Wang, K., Li, M., Hakonarson, H.: Analysing biological pathways in genome-wide association studies. Nature Reviews Genetics 11(12), 843–854 (2010)

[12] Yang, J., Ferreira, T., Morris, A.P., et al.: Conditional and joint multiple-SNP analysis of GWAS summary statistics identifies additional variants influencing complex traits. Nature Genetics 44(4), 369–375 (2012) 10.1038/ng.2213

[13] Wang, G., Sarkar, A., Carbonetto, P., Stephens, M.: A simple new approach to variable selection in regression, with application to genetic fine mapping. Journal of the Royal Statistical Society: Series B 82(5), 1273–1300 (2020) 10.1111/rssb.12388

[14] Benjamini, Y., Hochberg, Y.: Controlling the false discovery rate: A practical and powerful approach to multiple testing. Journal of the Royal Statistical Society: Series B 57(1), 289–300 (1995) 10.1111/j.2517-6161.1995.tb02031.x

[15] Candès, E., Fan, Y., Janson, L., Lv, J.: Panning for gold: Model-X knockoffs for high-dimensional controlled variable selection. Journal of the Royal Statistical Society: Series B 80(3), 551–577 (2018) 10.1111/rssb.12265

[16] Barber, R.F., Candès, E.J.: Controlling the false discovery rate via knockoffs. The Annals of Statistics 43(5), 2055–2085 (2015) 10.1214/15-AOS1337

[17] Sesia, M., Sabatti, C., Candès, E.J.: Gene hunting with hidden Markov model knockoffs. Biometrika 106(1), 1–18 (2019) 10.1093/biomet/asy033

[18] He, Z., Liu, L., Wang, C., et al.: Identification of putative causal loci in whole-genome sequencing data via knockoff statistics. Nature Communications 12, 3152 (2021) 10.1038/s41467-021-22889-4

[19] Spector, A., Fithian, W.: Asymptotically Optimal Knockoff Statistics via the Masked Likelihood Ratio. Preprint; originally posted in 2022 (2024). https://arxiv.org/abs/2212.08766

[20] Urbut, S.M., Wang, G., Carbonetto, P., Stephens, M.: Flexible statistical methods for estimating and testing effects in genomic studies with multiple conditions. Nature Genetics 51(1), 187–195 (2019) 10.1038/s41588-018-0268-8

