## Supplementary Methods S1-S10, Supplementary Figs.∼S1-S15, and Supplementary Tables S1, S5, and S7 for "LKOM: Longitudinal Knockoff with Masked Likelihood Ratio for FDR-Controlled Association Discovery"

#### S1. Canonical ordering and construction of the masked data

Let $\boldsymbol{g}_j$ and $\widetilde{\boldsymbol{g}}_j$ denote the original and knockoff genotype vectors for variant $j$ among the $m$ study participants. The marginal screening rule is applied to the pair as a whole: if either member passes the screening criterion in either response channel, both members are retained. Conditional on the retained pairs, LKOM removes the original–knockoff labels before fitting the Bayesian working model.

For each pair, a deterministic lexicographic rule maps the unordered pair $\{\boldsymbol{g}_j, \widetilde{\boldsymbol{g}}_j\}$ to a canonical ordered pair $(\boldsymbol{a}_j, \boldsymbol{b}_j)$. More precisely, let

$$d_j = \min\{i : |g_{ij} - \widetilde{g}_{ij}| > \delta\},$$

where $\delta \geq 0$ is a numerical tolerance. If $d_j$ exists, the member having the smaller value at position $d_j$ is assigned to $\boldsymbol{a}_j$ and the other member is assigned to $\boldsymbol{b}_j$. Thus, $\boldsymbol{a}_j$ is the lexicographically smaller of the two vectors. The analyses reported in this study used $\delta = 0$. If no such position exists, the two vectors are identical; the pair is flagged as non-informative and its final knockoff statistic is set to zero.

This construction is invariant to relabeling the two members of a pair. Indeed, exchanging $\boldsymbol{g}_j$ and $\widetilde{\boldsymbol{g}}_j$ leaves $(\boldsymbol{a}_j, \boldsymbol{b}_j)$, and hence $\mathcal{D}_{\text{mask}}$, unchanged, while reversing $o_j$. Therefore the canonical posterior log-odds is unchanged and the restored statistic changes sign. This property also holds when any subset of pairs is exchanged simultaneously.

#### S2. Complete likelihood and prior specification

Let $S_j \in \{A, B\}$ denote the latent orientation of pair $j$, with

$$\boldsymbol{x}_{j,S_j} = \begin{cases} \boldsymbol{a}_j, & S_j = A, \\ \boldsymbol{b}_j, & S_j = B. \end{cases}$$

Let $\boldsymbol{\theta}_j = (\theta_{\beta j}, \theta_{\tau j})^T$ be its coefficients for the mean and within-subject (WS) variability response channels. The dual-channel working likelihood is

$$\boldsymbol{Y} = \sum_{j=1}^{p} \boldsymbol{x}_{j,S_j} \boldsymbol{\theta}_j^T + \boldsymbol{E}, \qquad \boldsymbol{E}_{i\cdot}^T \overset{\text{iid}}{\sim} N_2(\boldsymbol{0}, \boldsymbol{\Omega}). \tag{1}$$

If

$$\boldsymbol{E}(\boldsymbol{\Theta}, \boldsymbol{S}) = \boldsymbol{Y} - \sum_{j=1}^{p} \boldsymbol{x}_{j,S_j} \boldsymbol{\theta}_j^T,$$

then

$$p(\boldsymbol{Y} \mid \boldsymbol{\Theta}, \boldsymbol{S}, \boldsymbol{\Omega}) = (2\pi)^{-m} |\boldsymbol{\Omega}|^{-m/2} \exp\left[-\frac{1}{2} \operatorname{tr}\left\{\boldsymbol{\Omega}^{-1} \boldsymbol{E}^T \boldsymbol{E}\right\}\right]. \tag{2}$$

Each orientation has prior probability 1/2. The association state $H_j \in \mathcal{H} = \{00, 10, 01, 11\}$ has probabilities $\boldsymbol{\pi} = (\pi_{00}, \pi_{10}, \pi_{01}, \pi_{11})^T$:

$$\Pr(S_j = A) = \Pr(S_j = B) = \frac{1}{2}, \qquad H_j \mid \boldsymbol{\pi} \overset{\text{iid}}{\sim} \text{Categorical}(\boldsymbol{\pi}), \qquad \boldsymbol{\pi} \sim \text{Dirichlet}(\kappa \boldsymbol{\pi}_0).$$

For the analyses reported here,

$$\boldsymbol{\pi}_0 = (0.90, 0.04, 0.04, 0.02)^T, \qquad \kappa = 20.$$

The active coordinate sets are $\mathcal{A}_{00} = \varnothing$, $\mathcal{A}_{10} = \{1\}$, $\mathcal{A}_{01} = \{2\}$, and $\mathcal{A}_{11} = \{1, 2\}$. Conditional on $H_j = h$, the inactive coefficients are fixed at zero and

$$\boldsymbol{\theta}_{j,\mathcal{A}_h} \mid H_j = h, s_\beta, s_\tau, \rho \sim N_{d_h}(\mathbf{0}, \boldsymbol{\Psi}_h), \qquad d_h = |\mathcal{A}_h|,$$

where

$$\boldsymbol{\Psi}_{10} = s_\beta^2, \qquad \boldsymbol{\Psi}_{01} = s_\tau^2, \qquad \boldsymbol{\Psi}_{11} = \begin{pmatrix} s_\beta^2 & \rho s_\beta s_\tau \\ \rho s_\beta s_\tau & s_\tau^2 \end{pmatrix}.$$

The residual covariance has the inverse-Wishart prior

$$\boldsymbol{\Omega} \sim \text{IW}_2(\nu_0, \boldsymbol{S}_0), \qquad \nu_0 = 5, \qquad \boldsymbol{S}_0 = 2\boldsymbol{I}_2, \tag{3}$$

under the parameterization

$$p(\boldsymbol{\Omega}) \propto |\boldsymbol{\Omega}|^{-(\nu_0+3)/2} \exp\left\{-\frac{1}{2}\operatorname{tr}(\boldsymbol{S}_0 \boldsymbol{\Omega}^{-1})\right\}.$$

Thus, $E(\boldsymbol{\Omega}) = \boldsymbol{I}_2$ under this prior, consistent with the standardized response scale.

The slab scales are supported on

$$\mathcal{G}_\beta = \mathcal{G}_\tau = \{0.02, 0.03, \ldots, 0.30\}.$$

For $k \in \{\beta, \tau\}$, the prior mass at $g \in \mathcal{G}_k$ is proportional to the density of a half-$t$ distribution with 3 degrees of freedom and scale 0.10 evaluated at $g$. The slab correlation is supported on

$$\mathcal{G}_\rho = \{-0.8, -0.7, \ldots, 0.8\},$$

with prior mass

$$\Pr(\rho = r) \propto (1 - r^2)^{\eta_\rho - 1}, \qquad \eta_\rho = 2,$$

which is the two-dimensional LKJ correlation prior evaluated on the grid. The priors for $s_\beta$, $s_\tau$, and $\rho$ are mutually independent before observing the data.

Let $n_h = \sum_{j=1}^{p} \mathbf{1}(H_j = h)$ and let $\varphi_{d_h}(\cdot; \mathbf{0}, \boldsymbol{\Psi}_h)$ denote the $d_h$-variate Gaussian density. Up to a normalizing constant, the complete posterior is

$$\begin{aligned} & p(\boldsymbol{\Theta}, \boldsymbol{S}, \boldsymbol{H}, \boldsymbol{\pi}, \boldsymbol{\Omega}, s_\beta, s_\tau, \rho \mid \mathcal{D}_{\text{mask}}) \\ & \quad \propto |\boldsymbol{\Omega}|^{-(m+\nu_0+3)/2} \exp\left[-\frac{1}{2}\operatorname{tr}\left\{(\boldsymbol{S}_0 + \boldsymbol{E}^T\boldsymbol{E})\boldsymbol{\Omega}^{-1}\right\}\right] \\ & \qquad \times 2^{-p} \prod_{h\in\mathcal{H}} \pi_h^{\kappa\pi_{0h}-1+n_h} \prod_{j:H_j\neq 00} \varphi_{d_{H_j}}\left(\boldsymbol{\theta}_{j,\mathcal{A}_{H_j}}; \mathbf{0}, \boldsymbol{\Psi}_{H_j}\right) \\ & \qquad \times p(s_\beta)p(s_\tau)p(\rho), \end{aligned} \tag{4}$$

with point masses at zero for all inactive coefficients. The Gaussian slab densities in Equation (4) include their determinant terms and therefore contribute to the full conditional updates of the slab scales and correlation.

#### S3. Derivation of the conditional Bayes factor

To update pair $j$, remove its current fitted contribution and define

$$\boldsymbol{R}_{-j} = \boldsymbol{Y} - \sum_{\ell\neq j} \boldsymbol{x}_{\ell,S_\ell}\boldsymbol{\theta}_\ell^T.$$

For a proposed orientation $s \in \{A, B\}$, write

$$q_{js} = \boldsymbol{x}_{j,s}^T\boldsymbol{x}_{j,s}, \qquad \boldsymbol{u}_{js} = \boldsymbol{R}_{-j}^T\boldsymbol{x}_{j,s}, \qquad \boldsymbol{\Lambda} = \boldsymbol{\Omega}^{-1}.$$

The part of the conditional log-likelihood that depends on $\boldsymbol{\theta}_j$ is

$$\boldsymbol{\theta}_j^T\boldsymbol{\Lambda}\boldsymbol{u}_{js} - \frac{1}{2}q_{js}\boldsymbol{\theta}_j^T\boldsymbol{\Lambda}\boldsymbol{\theta}_j. \tag{5}$$

For a non-null state $h$, restrict Equation (5) to the active coordinates and define

$$\boldsymbol{v}_{jsh} = (\boldsymbol{\Lambda}\boldsymbol{u}_{js})_{\mathcal{A}_h}, \qquad \boldsymbol{J}_{jsh} = q_{js}\boldsymbol{\Lambda}_{\mathcal{A}_h,\mathcal{A}_h} + \boldsymbol{\Psi}_h^{-1}.$$

Multiplication by the $N_{d_h}(\mathbf{0}, \boldsymbol{\Psi}_h)$ slab density gives the kernel

$$|\boldsymbol{\Psi}_h|^{-1/2} \exp\left\{-\frac{1}{2}\boldsymbol{\theta}_{j,\mathcal{A}_h}^T\boldsymbol{J}_{jsh}\boldsymbol{\theta}_{j,\mathcal{A}_h} + \boldsymbol{\theta}_{j,\mathcal{A}_h}^T\boldsymbol{v}_{jsh}\right\}.$$

Completing the square with $\boldsymbol{\mu}_{jsh} = \boldsymbol{J}_{jsh}^{-1}\boldsymbol{v}_{jsh}$ yields

$$-\frac{1}{2}(\boldsymbol{\theta}_{j,\mathcal{A}_h} - \boldsymbol{\mu}_{jsh})^T \boldsymbol{J}_{jsh}(\boldsymbol{\theta}_{j,\mathcal{A}_h} - \boldsymbol{\mu}_{jsh}) + \frac{1}{2}\boldsymbol{v}_{jsh}^T \boldsymbol{J}_{jsh}^{-1}\boldsymbol{v}_{jsh}.$$

Integrating over the active coefficients and dividing by the conditional likelihood under $H_j = 00$ therefore gives

$$\mathrm{BF}_{jsh} = |\boldsymbol{\Psi}_h|^{-1/2}|\boldsymbol{J}_{jsh}|^{-1/2} \exp\left(\frac{1}{2}\boldsymbol{v}_{jsh}^T \boldsymbol{J}_{jsh}^{-1}\boldsymbol{v}_{jsh}\right), \qquad h \neq 00, \tag{6}$$

with $\mathrm{BF}_{js,00} = 1$.

The eight orientation–state combinations have normalized conditional probabilities

$$p_{jsh} = \frac{\frac{1}{2}\pi_h \, \mathrm{BF}_{jsh}}{\displaystyle\sum_{s' \in \{A,B\}} \sum_{h' \in \mathcal{H}} \frac{1}{2}\pi_{h'} \, \mathrm{BF}_{js'h'}}. \tag{7}$$

Conditional on selecting $(S_j, H_j) = (s, h)$, the active effects follow

$$\boldsymbol{\theta}_{j,\mathcal{A}_h} \mid \cdot \sim N_{d_h}\left(\boldsymbol{J}_{jsh}^{-1}\boldsymbol{v}_{jsh}, \boldsymbol{J}_{jsh}^{-1}\right), \tag{8}$$

and all inactive coordinates are set to zero.

#### S4. Gibbs sampler and posterior computation

The implementation stores both members of every canonical pair in

$$\boldsymbol{X} = [\boldsymbol{A}, \boldsymbol{B}] \in \mathbb{R}^{m \times 2p}$$

and uses a coefficient matrix $\boldsymbol{C} \in \mathbb{R}^{2p \times 2}$. For each pair, at most one of rows $j$ and $p + j$ is nonzero. Thus, $\boldsymbol{X}\boldsymbol{C} = \sum_j \boldsymbol{x}_{j,S_j}\boldsymbol{\theta}_j^T$. The matrices

$$\boldsymbol{Q} = \boldsymbol{X}^T\boldsymbol{X}, \qquad \boldsymbol{T} = \boldsymbol{X}^T\boldsymbol{Y}$$

are computed once. If $I_j = \{j, p + j\}$, the two score vectors required for the update of pair $j$ are evaluated without reconstructing the full residual:

$$\begin{pmatrix} \boldsymbol{u}_{jA}^T \\ \boldsymbol{u}_{jB}^T \end{pmatrix} = \boldsymbol{T}_{I_j,\cdot} - \boldsymbol{Q}_{I_j,\cdot}\boldsymbol{C} + \boldsymbol{Q}_{I_j,I_j}\boldsymbol{C}_{I_j,\cdot}. \tag{9}$$

One systematic Gibbs sweep consists of the following updates.

1. For $j = 1, \ldots, p$, calculate the eight probabilities in Equation (7), sample $(S_j, H_j)$, draw the active effects from Equation (8), and replace the two corresponding rows of $\boldsymbol{C}$. The update uses the most recent values of pairs $1, \ldots, j-1$ and the preceding-sweep values of the remaining pairs.
2. Given the updated state counts, sample

$$\boldsymbol{\pi} \mid \boldsymbol{H} \sim \text{Dirichlet}\left(n_{00} + \kappa\pi_{0,00}, n_{10} + \kappa\pi_{0,10}, n_{01} + \kappa\pi_{0,01}, n_{11} + \kappa\pi_{0,11}\right).$$

3. Let $\boldsymbol{E} = \boldsymbol{Y} - \boldsymbol{X}\boldsymbol{C}$. Sample

$$\boldsymbol{\Omega} \mid \cdot \sim \text{IW}_2\left(\nu_0 + m, \boldsymbol{S}_0 + \boldsymbol{E}^T\boldsymbol{E}\right).$$

4. Update $s_\beta$, $s_\tau$, and $\rho$ sequentially by enumerating their finite grids. For example,

$$\Pr(s_\beta = g \mid \cdot) \propto p(s_\beta = g) \prod_{j: H_j \neq 00} \varphi_{d_{H_j}}\left(\boldsymbol{\theta}_{j,\mathcal{A}_{H_j}}; \boldsymbol{0}, \boldsymbol{\Psi}_{H_j}(g, s_\tau, \rho)\right),$$

and analogous expressions are used for $s_\tau$ and $\rho$. Probabilities are normalized on the relevant grid and one value is sampled. This update retains both the quadratic and determinant contributions of the slab density, including the correlation term for state 11.

The working covariance is initialized as

$$\boldsymbol{\Omega}^{(0)} = m^{-1}\boldsymbol{Y}^T\boldsymbol{Y} + 10^{-6}\boldsymbol{I}_2,$$

and the initial slab values are $s_\beta = s_\tau = 0.10$ and $\rho = 0$. The Gibbs sampler was initialized using the deterministic conditional-Bayes-factor procedure described in Supplementary Method 5. Each sweep updated the variant pairs in a fixed forward order. For the reported simulations, 100 burn-in sweeps were followed by 300 retained sweeps without thinning.

At each retained pair update, LKOM records the conditional probabilities $p_{jsh}$ rather than only the sampled state. With $M$ retained sweeps, the Rao–Blackwellized posterior summaries are

$$\widehat{P}_{jA} = \frac{1}{M}\sum_{t=1}^{M}\sum_{h\in\mathcal{H}} p_{jAh}^{(t)}, \qquad \widehat{P}_{jh} = \frac{1}{M}\sum_{t=1}^{M}\left(p_{jAh}^{(t)} + p_{jBh}^{(t)}\right).$$

#### S5. Deterministic conditional-Bayes-factor initialization

LKOM provides a deterministic conditional-Bayes-factor (conBF) procedure to initialize the Gibbs sampler. This initialization aims to place the chain in a region of relatively high posterior support and improve the stability of subsequent sampling. The procedure uses the conditional probabilities derived in Supplementary

Method 3 to obtain initial effect coefficients, association states, and state proportions. The residual covariance $\boldsymbol{\Omega}$ and slab parameters $(s_\beta, s_\tau, \rho)$ remain fixed at their initial values during this procedure.

**Coordinate updates.** Using the augmented canonical design matrix $\boldsymbol{X} = [\boldsymbol{A}, \boldsymbol{B}]$ and coefficient matrix $\boldsymbol{C}$ defined in Supplementary Method 4, each conBF fit starts from $\boldsymbol{C}^{(0)} = \boldsymbol{0}$ and $\boldsymbol{\pi}^{(0)} = \boldsymbol{\pi}_0$. For each pair $j$, the partial-residual scores are calculated using the precomputed matrices $\boldsymbol{Q} = \boldsymbol{X}^T\boldsymbol{X}$ and $\boldsymbol{T} = \boldsymbol{X}^T\boldsymbol{Y}$, as in Equation (9). The conditional Bayes factors then give the probabilities

$$p_{jsh} = \frac{\frac{1}{2}\pi_h \,\mathrm{BF}_{jsh}}{\displaystyle\sum_{s' \in \{A,B\}} \sum_{h' \in \mathcal{H}} \frac{1}{2}\pi_{h'} \,\mathrm{BF}_{js'h'}}.$$

Instead of sampling an orientation–state combination, conBF selects

$$(\widehat{S}_j, \widehat{H}_j) = \operatorname*{arg\,max}_{s \in \{A,B\},\, h \in \mathcal{H}} p_{jsh}.$$

For a non-null state, the active coefficients are replaced by their conditional posterior mean,

$$\widehat{\boldsymbol{\theta}}_{j,\mathcal{A}_{\widehat{H}_j}} = \boldsymbol{J}^{-1}_{j,\widehat{S}_j,\widehat{H}_j} \boldsymbol{v}_{j,\widehat{S}_j,\widehat{H}_j},$$

while inactive coefficients are set to zero. If $\widehat{H}_j = 00$, the entire effect vector is set to zero. Only the row of $\boldsymbol{C}$ corresponding to $\widehat{S}_j$ receives the updated effect vector; the other row of the pair is set to zero. Pairs are updated sequentially, so each update uses the most recent coefficient estimates for previously visited pairs.

The state proportions are held fixed within each sweep. After completing a sweep, they are updated using the soft state probabilities:

$$\pi_h^{(t+1)} = \frac{\displaystyle\sum_{j=1}^{p} \sum_{s \in \{A,B\}} p_{jsh}^{(t)} + \kappa \pi_{0h}}{p + \kappa}, \qquad h \in \mathcal{H},$$

where $p_{jsh}^{(t)}$ denotes the conditional probability calculated when pair $j$ was visited during sweep $t$. The shrinkage center $\boldsymbol{\pi}_0$ remains fixed throughout the fit.

Iteration stops when

$$\Delta^{(t)} = \max_{a,b} \left| C_{ab}^{(t+1)} - C_{ab}^{(t)} \right| \le 10^{-6},$$

or after 100 sweeps, whichever occurs first. This stopping rule measures stabilization of the coefficient estimates. After the final sweep, the conditional orientation and state probabilities are recalculated for every pair using the final coefficient matrix and state proportions.

#### S6. Swap invariance, antisymmetry, and FDR control

Let $\boldsymbol{Z} = [\boldsymbol{G}, \widetilde{\boldsymbol{G}}]$ denote the augmented genotype matrix, and let $\boldsymbol{Z}_{\mathrm{swap}(\mathcal{S})}$ be obtained by exchanging $\boldsymbol{G}_j$ and $\widetilde{\boldsymbol{G}}_j$ for every $j \in \mathcal{S}$, where $\mathcal{S} \subseteq \{1, \ldots, p\}$. This section establishes the flip-sign property of the LKOM statistics and the resulting false discovery rate control for the dual-channel joint null.

**Proposition 1 (Flip-sign property).** For any $\mathcal{S} \subseteq \{1, \ldots, p\}$, the LKOM statistics satisfy

$$W_j\big(\boldsymbol{Z}_{\mathrm{swap}(\mathcal{S})}, \boldsymbol{Y}\big) = \begin{cases} -W_j(\boldsymbol{Z}, \boldsymbol{Y}), & j \in \mathcal{S}, \\ W_j(\boldsymbol{Z}, \boldsymbol{Y}), & j \notin \mathcal{S}. \end{cases} \tag{10}$$

*Proof.* We establish the result by considering each step of the LKOM construction.

**Invariance of the retained pair set.** The marginal screening rule retains pair $j$ according to a symmetric function of the association evidence for its two members. For example, in the dual-channel analysis, the pair is retained when

$$\min\left(p_{\beta j}, p_{\tau j}, \widetilde{p}_{\beta j}, \widetilde{p}_{\tau j}\right) < p_{\mathrm{scr}}.$$

Exchanging the original and knockoff members only permutes the four arguments of this minimum and therefore leaves the retained pair set unchanged. Variants excluded by screening are assigned $W_j = 0$, which also satisfies Equation (10).

**Invariance of the masked representation.** Let $\mathcal{C}(\boldsymbol{u}, \boldsymbol{v}) = (\boldsymbol{a}, \boldsymbol{b})$ denote the canonical-ordering rule defined in Supplementary Method 1. Because this rule depends only on the unordered pair $\{\boldsymbol{u}, \boldsymbol{v}\}$, $\mathcal{C}(\boldsymbol{g}_j, \widetilde{\boldsymbol{g}}_j) = \mathcal{C}(\widetilde{\boldsymbol{g}}_j, \boldsymbol{g}_j)$. Consequently, exchanging the observed labels leaves the canonical pair $(\boldsymbol{a}_j, \boldsymbol{b}_j)$ unchanged. In related samples, the whitening parameter is estimated from $\boldsymbol{Y}$ and the fixed GRM $\boldsymbol{\Phi}$, without using the observed original–knockoff labels. Moreover, the same linear transformation is applied to the two members of every pair. As shown in Supplementary Method 8, whitening commutes with paired column exchanges. Therefore, the complete masked data supplied to the Bayesian working model satisfy

$$\mathcal{D}_{\mathrm{mask}}\left(\boldsymbol{Z}_{\mathrm{swap}(\mathcal{S})}, \boldsymbol{Y}\right) = \mathcal{D}_{\mathrm{mask}}(\boldsymbol{Z}, \boldsymbol{Y}). \tag{11}$$

The observed orientation, by contrast, changes according to

$$o_j^{\mathrm{swap}(\mathcal{S})} = \begin{cases} -o_j, & j \in \mathcal{S}, \\ o_j, & j \notin \mathcal{S}. \end{cases} \tag{12}$$

If the original and knockoff genotype vectors are identical, LKOM sets $W_j = 0$, and the flip-sign identity remains valid because $-W_j = W_j = 0$.

**Invariance of posterior inference.** The conditional Bayes factors, Gibbs transition probabilities, and Rao–Blackwellized posterior summaries are functions of $\mathcal{D}_{\text{mask}}$ and the algorithmic random numbers used during posterior computation. Any such random numbers are generated independently of the observed original–knockoff labels. Conditional on a fixed realization of this external randomness, the same random-number stream is used for the original and swapped representations.

It follows from Equation (11) that

$$\widehat{P}_{jA}^{\text{swap}(\mathcal{S})} = \widehat{P}_{jA}, \qquad \widehat{P}_{jh}^{\text{swap}(\mathcal{S})} = \widehat{P}_{jh}, \quad h \in \mathcal{H}.$$

Therefore, the canonical orientation log-odds

$$L_j = \log\left\{\frac{\widehat{P}_{jA}}{1-\widehat{P}_{jA}}\right\}$$

is unchanged by exchanging any subset of the observed original–knockoff labels. Combining this invariance with Equation (12) gives

$$W_j^{\text{swap}(\mathcal{S})} = o_j^{\text{swap}(\mathcal{S})} L_j^{\text{swap}(\mathcal{S})} = \begin{cases} -o_j L_j, & j \in \mathcal{S}, \\ o_j L_j, & j \notin \mathcal{S}, \end{cases} = \begin{cases} -W_j, & j \in \mathcal{S}, \\ W_j, & j \notin \mathcal{S}. \end{cases}$$

This proves Equation (10). □

**Joint-null hypothesis.** For variant $j$, define the dual-channel joint null as

$$H_{0j} : \boldsymbol{Y} \perp\!\!\!\perp \boldsymbol{G}_j \mid \boldsymbol{G}_{-j}. \tag{13}$$

In the longitudinal location–scale model, this corresponds to the absence of both a mean effect and a WS-variability effect. Let $\mathcal{H}_0 = \{j : H_{0j} \text{ is true}\}$ denote the set of joint-null variants.

**Theorem 1 (FDR control of LKOM).** Suppose that the following conditions hold:

1. The knockoff genotypes satisfy model-X pairwise exchangeability,

$$(\boldsymbol{G}, \widetilde{\boldsymbol{G}})_{\text{swap}(\mathcal{S})} \overset{d}{=} (\boldsymbol{G}, \widetilde{\boldsymbol{G}}), \qquad \text{for every } \mathcal{S} \subseteq \{1, \ldots, p\}. \tag{14}$$

2. The knockoff genotypes satisfy conditional independence,

$$\widetilde{\boldsymbol{G}} \perp\!\!\!\perp \boldsymbol{Y} \mid \boldsymbol{G}. \tag{15}$$

3. Marginal screening and all preprocessing operations applied before construction of the LKOM statistics are invariant to exchanges of original–knockoff pairs.
4. Any algorithmic randomness used to construct the statistics is independent of the observed original–knockoff labels.

Let $\mathcal{T} = \{|W_j| : W_j \neq 0\}$ denote the set of candidate thresholds. For a target FDR level $q \in (0, 1)$, define the knockoff+ threshold as

$$T_q = \min\left\{t \in \mathcal{T} : \frac{1 + \#\{j : W_j \leq -t\}}{\#\{j : W_j \geq t\} \vee 1} \leq q\right\}, \tag{16}$$

and let $\widehat{\mathcal{S}}_q = \{j : W_j \geq T_q\}$. If the set in Equation (16) is empty, we set $T_q = \infty$ and $\widehat{\mathcal{S}}_q = \varnothing$. Then

$$\mathrm{FDR} = \mathbb{E}\left[\frac{|\widehat{\mathcal{S}}_q \cap \mathcal{H}_0|}{|\widehat{\mathcal{S}}_q| \vee 1}\right] \leq q. \tag{17}$$

*Proof.* By Proposition 1, the LKOM statistics satisfy the flip-sign property under the exchange of any subset of original–knockoff pairs. Under Equations (14) and (15), exchanging any subset of joint-null original–knockoff pairs leaves the joint distribution of the augmented genotypes and response unchanged.

Consequently, conditional on the statistic magnitudes and the statistics for non-null variants, the signs of the nonzero statistics corresponding to joint-null variants are independent Rademacher variables. In particular, a sufficiently large negative null statistic is as likely as a positive null statistic of the same magnitude. The numerator of Equation (16), including the offset of one, therefore provides a conservative estimate of the number of false discoveries among the statistics exceeding the positive threshold. □

Thus, LKOM controls variant-level FDR with respect to the joint null of no association with either response channel. The validity of this result depends on the model-X knockoff assumptions and the swap symmetry of the complete statistic-construction procedure. The four-state posterior probabilities are used to characterize the association patterns of selected variants and do not enter the knockoff+ threshold.

#### S7. Profile REML estimation and GRM-based whitening

Let $\boldsymbol{\Phi}$ be the standardized genetic relationship matrix (GRM), aligned with the $m$ participants, and let $\boldsymbol{Y} \in \mathbb{R}^{m\times k}$ denote the working response, with $k = 2$ for dual-channel LKOM. The null relatedness model is

$$\boldsymbol{Y} = \mathbf{1}_m\boldsymbol{\mu}^T + \boldsymbol{E}, \qquad \boldsymbol{E} \sim \mathrm{MN}_{m\times k}(\mathbf{0}, \boldsymbol{V}_\eta, \boldsymbol{\Sigma}_R), \qquad \boldsymbol{V}_\eta = (1-\eta)\boldsymbol{I}_m + \eta\boldsymbol{\Phi}. \tag{18}$$

For a fixed $\eta$, define

$$c_\eta = \mathbf{1}_m^T \boldsymbol{V}_\eta^{-1} \mathbf{1}_m, \qquad \widehat{\boldsymbol{\mu}}(\eta)^T = \frac{\mathbf{1}_m^T \boldsymbol{V}_\eta^{-1} \boldsymbol{Y}}{c_\eta},$$

and

$$\boldsymbol{E}_\eta = \boldsymbol{Y} - \mathbf{1}_m \widehat{\boldsymbol{\mu}}(\eta)^T, \qquad \widehat{\boldsymbol{\Sigma}}_R(\eta) = \frac{1}{m-1} \boldsymbol{E}_\eta^T \boldsymbol{V}_\eta^{-1} \boldsymbol{E}_\eta.$$

Up to an additive constant, the restricted negative log-likelihood before profiling out $\boldsymbol{\Sigma}_R$ is

$$\begin{aligned} \mathcal{L}_R(\eta, \boldsymbol{\Sigma}_R) = &\frac{k}{2} \log |\boldsymbol{V}_\eta| + \frac{k}{2} \log c_\eta + \frac{m-1}{2} \log |\boldsymbol{\Sigma}_R| \\ &+ \frac{1}{2} \operatorname{tr}\left(\boldsymbol{\Sigma}_R^{-1} \boldsymbol{E}_\eta^T \boldsymbol{V}_\eta^{-1} \boldsymbol{E}_\eta\right). \end{aligned} \tag{19}$$

Differentiation with respect to $\boldsymbol{\Sigma}_R$ gives the estimator above. Substitution into Equation (19) yields

$$\mathcal{L}_{R,p}(\eta) = \frac{k}{2} \{\log |\boldsymbol{V}_\eta| + \log c_\eta\} + \frac{m-1}{2} \log |\widehat{\boldsymbol{\Sigma}}_R(\eta)|. \tag{20}$$

For $k = 2$, this reduces to

$$\mathcal{L}_{R,p}(\eta) = \log |\boldsymbol{V}_\eta| + \log c_\eta + \frac{m-1}{2} \log |\widehat{\boldsymbol{\Sigma}}_R(\eta)|,$$

which is the objective used for dual-channel LKOM. The implementation minimizes this one-dimensional objective over $[0, 0.999]$, with numerical tolerance $10^{-6}$, and explicitly compares the interior optimizer with both endpoints.

If $\boldsymbol{\Phi} = \boldsymbol{U} \operatorname{diag}(\lambda_1, \ldots, \lambda_m) \boldsymbol{U}^T$, then

$$\begin{aligned} \boldsymbol{V}_\eta^{-1} &= \boldsymbol{U} \operatorname{diag}\left\{(1 - \eta + \eta\lambda_i)^{-1}\right\} \boldsymbol{U}^T, \\ \boldsymbol{V}_\eta^{-1/2} &= \boldsymbol{U} \operatorname{diag}\left\{(1 - \eta + \eta\lambda_i)^{-1/2}\right\} \boldsymbol{U}^T. \end{aligned}$$

In the pedigree simulations, the sparse GRM was block diagonal by family. The implementation therefore diagonalized each family block separately, which is algebraically identical to diagonalizing the full block-diagonal matrix and avoids forming a dense $m \times m$ GRM.

After estimating $\widehat{\eta}$ from the response and GRM alone, define the linear transformation

$$\mathcal{T}_{\widehat{\eta}}(\boldsymbol{D}) = \boldsymbol{V}_{\widehat{\eta}}^{-1/2} \left[\boldsymbol{D} - \mathbf{1}_m \frac{\mathbf{1}_m^T \boldsymbol{V}_{\widehat{\eta}}^{-1} \boldsymbol{D}}{\mathbf{1}_m^T \boldsymbol{V}_{\widehat{\eta}}^{-1} \mathbf{1}_m}\right]. \tag{21}$$

Generalized least-squares centering is performed before multiplication by $\boldsymbol{V}_{\widehat{\eta}}^{-1/2}$; ordinary centering after whitening is not equivalent unless $\boldsymbol{V}_{\widehat{\eta}}^{-1/2} \mathbf{1}_m$ is constant. LKOM applies Equation (21) to $\boldsymbol{Y}$, $\boldsymbol{G}$, and $\widetilde{\boldsymbol{G}}$, after which the transformed columns are standardized as described in Section S1. Dual-channel

LKOM estimates one $\widehat{\eta}$ from the bivariate response. For a single-channel analysis, the same profile REML calculation is applied with $k = 1$ to the corresponding response channel.

Finally, whitening is equivariant to original–knockoff exchanges. Let $\boldsymbol{Z} = [\boldsymbol{G}, \widetilde{\boldsymbol{G}}]$ and let $\boldsymbol{P}_{\mathcal{S}}$ exchange any subset $\mathcal{S}$ of paired columns. Because $\widehat{\eta}$ depends only on $\boldsymbol{Y}$ and $\boldsymbol{\Phi}$, and because the same row transformation is applied to every genotype column, $\mathcal{T}_{\widehat{\eta}}(\boldsymbol{Z}\boldsymbol{P}_{\mathcal{S}}) = \mathcal{T}_{\widehat{\eta}}(\boldsymbol{Z})\boldsymbol{P}_{\mathcal{S}}$. Thus, swapping a genotype with its knockoff commutes with both GLS centering and whitening. The masked canonical data remain unchanged under the swap, whereas the observed orientation changes sign, preserving the antisymmetry of the LKOM statistic.

#### S8. UK Biobank genotype preparation and knockoff generation

**Selection of unrelated participants.** The genotype data used in the primary simulations were derived from quality-controlled imputed genotypes in White British participants from UK Biobank. To define an unrelated source population, we first selected autosomal, biallelic A/C/G/T variants with minor allele frequency (MAF) at least 0.05, variant and sample missingness no greater than 0.02, and Hardy–Weinberg equilibrium test $p \geq 10^{-6}$. These variants were LD pruned using a window of 200 variants, a step size of 50 variants, and a pairwise $r^2$ threshold of 0.1.

Pairwise relatedness was then evaluated using KING as implemented in PLINK 2.0. A kinship cutoff of 0.0442 was used so that no two participants in the retained set were third-degree relatives or closer. Participant identifiers in the resulting maximal unrelated set were sorted, and 10,000 participants were sampled without replacement using random seed 718. The same set of participants was used throughout the primary simulations.

**Genotype processing.** Genotypes were extracted for the selected participants. The upstream genotype resource had already been filtered at MAF $\geq 2 \times 10^{-4}$. Within the 10,000-participant analysis cohort, variants were additionally required to have minor allele count (MAC) at least 25, genotype missingness no greater than 0.02, and Hardy–Weinberg equilibrium test $p \geq 10^{-6}$. At this sample size, the MAC requirement corresponds to a minimum possible MAF of $25/2 \times 10{,}000 = 1.25 \times 10^{-3}$, and was therefore more restrictive than the upstream MAF threshold.

To reduce very high correlations between original and knockoff variables, variants were LD pruned using `--indep-pairwise 10000kb 1 0.75`. The retained variants were assigned to the genomic intervals specified in the LD-block map. For each simulation replicate, one starting block was sampled uniformly from the allowable positions, and five physically consecutive.

**Knockoff construction.** Knockoff genotypes were generated using `create.MK` in the R package `KnockoffScreen` (version 0.3.0). Consecutive LD blocks were processed jointly and the resulting knockoff matrix was subsequently divided into block-specific files while preserving the original SNP and participant order. One knockoff copy was generated for each variant. The principal `create.MK` settings were

$$M = 1, \qquad \texttt{corr\_max} = 0.75, \qquad \texttt{maxN.neighbor} = \infty,$$

$$\texttt{maxBP.neighbor} = 100{,}000, \qquad \texttt{thres.ultrarare} = 25, \qquad \texttt{R2.thres} = 1.$$

The shrinkage construction was used for groups containing more than one variant. Genotypes missing within the input to `create.MK` were mean imputed by the genotype-reading routine.

#### S9. Detailed longitudinal phenotype generation

For subject $i = 1, \ldots, m$, the number of longitudinal measurements was sampled independently from the discrete uniform distribution

$$n_i \sim \text{Unif}\{6, \ldots, 15\}.$$

Let $j = 1, \ldots, n_i$ index the measurements for subject $i$. The fixed-effect and random-effect covariates were specified as

$$\boldsymbol{x}_{ij} = \boldsymbol{w}_{ij} = (1, x_{1i}, x_{2i}, x_{3ij})^T, \qquad \boldsymbol{z}_{ij} = (1, z_{ij})^T.$$

The subject-level covariates were generated as

$$x_{1i} \sim \text{Bernoulli}(0.5), \qquad x_{2i} \sim N(0, 0.5^2),$$

and the measurement-level covariates were generated independently as

$$x_{3ij} \sim N(0, 0.3^2), \qquad z_{ij} \sim N(0, 0.5^2).$$

We set

$$\boldsymbol{\beta} = (1, 0.5, 0.5, -0.3)^T, \qquad \boldsymbol{\tau} = (0.25, 0.30, -0.15, 0.10)^T.$$

The subject-specific mean and variability random effects were generated according to

$$(\gamma_{0i}, \gamma_{1i}, \omega_i)^T \overset{\text{ind}}{\sim} N(\mathbf{0}, 0.5\boldsymbol{I}_3),$$

so that the random contribution to the mean trajectory was

$$\boldsymbol{z}_{ij}^T \gamma_i = \gamma_{0i} + z_{ij}\gamma_{1i}.$$

**Selection of causal variants.** Let $\mathcal{C}_\beta$ and $\mathcal{C}_\tau$ denote the causal variant sets for the mean and WS variability channels, respectively. For each channel, causal-source blocks were sampled from the LD blocks included in the replicate. Under the default setting, five causal-source blocks were used, so that variants in all five analysis blocks were eligible to be causal.

Causal variants were sampled without replacement from the corresponding candidate pools. The default candidate pool included all eligible variants. In MAF-stratified simulations, the pool was restricted to rare variants with MAF $< 0.01$, low-frequency variants with $0.01 \leq$ MAF $< 0.05$, or common variants with MAF $\geq 0.05$.

When a positive overlap proportion $r_{\text{ov}}$ was specified, the target number of variants shared between the two channels was

$$n_{\text{shared}} = \lfloor r_{\text{ov}} \min\left(|\mathcal{C}_\beta|, |\mathcal{C}_\tau|\right) + 0.5 \rfloor .$$

The remaining causal variants were sampled separately for the two channels, subject to the constraint that the realized number of shared variants equaled $n_{\text{shared}}$. When $r_{\text{ov}} = 0$, the two causal sets were sampled independently without imposing disjointness; incidental overlap could therefore occur by chance.

**MAF-dependent genetic effects.** For channel $c \in \{\beta, \tau\}$ and causal variant $k \in \mathcal{C}_c$, an initial effect was generated as

$$a_{ck} = -\log_{10}(\text{MAF}_k) u_{ck}, \qquad u_{ck} \overset{\text{ind}}{\sim} \text{Unif}(0.2, 0.5).$$

The initial effects were positive under the genotype dosage coding used in the simulations; their signs were not independently randomized. For a variant affecting both channels, the effect multipliers were generated independently for the two channels.

Using the raw, unstandardized allele dosages $G_{ik}$, the initial subject-level genetic scores were

$$S_{\beta i}^{(0)} = \sum_{k \in \mathcal{C}_\beta} G_{ik} a_{\beta k}, \qquad S_{\tau i}^{(0)} = \sum_{k \in \mathcal{C}_\tau} G_{ik} a_{\tau k}.$$

**Calibration of WS-variability effects.** The initial WS-variability score was rescaled to attain the specified $\text{PVE}_\tau$, defined on the subject-level log-variance predictor as

$$\text{PVE}_\tau = \frac{\text{Var}(S_\tau)}{\text{Var}(S_\tau) + \text{Var}(\omega)}.$$

Specifically, we set

$$c_\tau = \left\{ \frac{\text{PVE}_\tau}{1 - \text{PVE}_\tau} \frac{\text{Var}(\omega)}{\text{Var}(S_\tau^{(0)})} \right\}^{1/2}, \qquad S_{\tau i} = c_\tau S_{\tau i}^{(0)}.$$

The final effect of causal variant $k$ in the WS variability channel was therefore $c_\tau a_{\tau k}$.

The visit-specific log residual variance was then generated as

$$\eta_{\tau,ij} = \boldsymbol{w}_{ij}^T \boldsymbol{\tau} + S_{\tau i} + \omega_i,$$

with

$$\sigma_{ij}^2 = \exp(\eta_{\tau,ij}).$$

In the primary simulations,

$$e_{ij} \overset{\text{ind}}{\sim} N(0,1), \qquad \varepsilon_{ij} = \sigma_{ij} e_{ij}.$$

Thus, $\text{Var}(\varepsilon_{ij} \mid \boldsymbol{w}_{ij}, S_{\tau i}, \omega_i) = \sigma_{ij}^2$.

**Calibration of mean effects.** Because the mean-channel genetic score was defined at the subject level, $S_{\beta i}^{(0)}$ was repeated over the observed measurements for subject $i$ before calculating its empirical variance. Let

$$R_{ij}^{(0)} = \boldsymbol{z}_{ij}^T \boldsymbol{\gamma}_i + \varepsilon_{ij}$$

denote the non-genetic random contribution to the mean outcome. The mean-channel scaling factor was

$$c_\beta = \left\{ \frac{\text{PVE}_\beta}{1 - \text{PVE}_\beta} \frac{\text{Var}(R^{(0)})}{\text{Var}(S_\beta^{(0)})} \right\}^{1/2}, \qquad S_{\beta i} = c_\beta S_{\beta i}^{(0)},$$

where both variances were evaluated empirically over the longitudinal observations. Equivalently,

$$\text{PVE}_\beta = \frac{\text{Var}(S_\beta)}{\text{Var}(S_\beta) + \text{Var}\left(\boldsymbol{Z}\boldsymbol{\gamma} + \boldsymbol{\varepsilon}\right)}.$$

The final mean effect of causal variant $k$ was $c_\beta a_{\beta k}$.

Finally, the longitudinal phenotype was generated as

$$Y_{ij} = \boldsymbol{x}_{ij}^T \boldsymbol{\beta} + S_{\beta i} + \boldsymbol{z}_{ij}^T \boldsymbol{\gamma}_i + \varepsilon_{ij}.$$

If a channel contained no causal variants, its genetic score was set to zero. The use of replicate-specific empirical variances in the scaling factors maintained the target channel-specific PVE when the number, MAF distribution, or overlap of causal variants was changed.

For the heavy-tailed residual sensitivity analysis, the standard normal innovation $e_{ij}$ was replaced by

$$e_{ij} = t_{5,ij} \sqrt{\frac{5-2}{5}},$$

which has unit variance. For the reduced-follow-up sensitivity analysis, $n_i$ was instead sampled uniformly from $\{2, \ldots, 8\}$; all other phenotype-generation parameters were unchanged.

#### S10. Related-sample simulation designs

We considered two related-sample designs. The first used regular pedigrees generated by GRAB, whereas the second preserved the sparse and irregular relatedness structure observed among UK Biobank participants. Each simulated cohort contained 10,000 individuals.

**GRAB-generated pedigree cohorts.** Family-structured genotypes were generated from the UK Biobank-derived genotypes using `GRAB.SimuGMatFromGenoFile` in the R package `GRAB` (version 0.2.4). We considered three family-size configurations:

| Family size | Number of families | Number of unrelated individuals |
|---|---|---|
| 4 | 1,250 | 5,000 |
| 10 | 500 | 5,000 |
| 20 | 250 | 5,000 |

so that the members of the simulated families and the unrelated individuals each contributed 5,000 participants.

After genotype generation, variants with MAC below 25 were removed. Variant identifiers, physical positions, reference alleles, alternative alleles, and dosage directions were restored from the source UK Biobank marker information before writing the simulated genotypes to PLINK format. Genotype and marker dimensions were checked after each conversion. The simulated genotypes were divided according to the same LD-block map used in the primary simulations, and knockoffs were generated separately for each family-size configuration using the procedure described in Supplementary Method 9.

A sparse GRM was constructed within each simulated cohort using the GRAB interface to GCTA. Off-diagonal entries linking individuals from different simulated families were set to zero, and diagonal entries were set to one. Pairwise IBD quantities required by the related-sample comparison methods were calculated from the same cohort-specific genotypes.

**UK Biobank cohorts with observed relatedness.** The second design sampled participants directly from UK Biobank. A participant was classified as related within a cohort if at least one participant with a kinship coefficient of at least 0.04419417 was also present in that cohort. This threshold corresponds to third-degree relatives or closer. We constructed three cohorts in which exactly 25%, 50%, or 75% of the 10,000 participants satisfied this within-cohort definition.

The kinship pairs were represented as an undirected graph, with participants as vertices and qualifying kinship pairs as edges. Disjoint anchor pairs were selected using a randomized greedy matching algorithm that preferentially considered edges whose two endpoints had low combined graph degree. Prioritizing low-degree pairs reduced the probability that additional unselected relatives would alter the target number of related participants.

The three cohorts were constructed jointly from these common related and filler pools. For a target related proportion $r$, the cohort contained $n_{\mathrm{rel}} = 10{,}000r$ participants from the related pool and $n_{\mathrm{fill}} = 10{,}000 - n_{\mathrm{rel}}$ participants from the filler pool. Selection used random seed 718. Each final cohort was explicitly checked to contain 10,000 unique participants and exactly 2,500, 5,000, or 7,500 participants with at least one third-degree-or-closer relative in the same cohort.

For GRM construction, we used an approximately 69,000-variant set of common, LD-pruned autosomal SNPs. Genotypes for the selected 10,000 participants were extracted separately for each cohort. The GRM calculation required MAF at least 0.01 and genotype missingness no greater than 0.10, and was divided into 20 computational parts before final assembly. The sparse GRM retained pairwise entries at or above the third-degree kinship threshold 0.04419417. Pairwise IBD information was additionally calculated.

**Generation of relatedness-induced phenotype effects.** Let $\mathbf{\Phi}$ denote the cohort-specific sparse GRM in the same participant order as the genotype matrices. Independently of the random effects defined in Supplementary Method 9, we generated

$$\boldsymbol{b}^{\beta} \sim N\left(\mathbf{0}, 0.3^2\mathbf{\Phi}\right), \qquad \boldsymbol{b}^{\tau} \sim N\left(\mathbf{0}, 0.2^2\mathbf{\Phi}\right).$$

The first vector induced relatedness in the trajectory mean, whereas the second induced relatedness in the log residual variance.

To avoid constructing a dense $10{,}000 \times 10{,}000$ matrix, the connected components of the sparse GRM were identified from its nonzero off-diagonal entries. Each component was diagonalized separately, and the Gaussian effects were generated from the resulting eigenvalues and eigenvectors. Small negative eigenvalues arising from numerical imprecision were truncated to zero. The same repaired component-wise eigensystem was subsequently used for GRM-based whitening.

For the related-sample simulations, the log-variance predictor was

$$\eta_{\tau,ij} = \boldsymbol{w}_{ij}^T\boldsymbol{\tau} + S_{\tau i} + \omega_i + b_i^{\tau},$$
$$\varepsilon_{ij} \sim N\left(0, \exp(\eta_{\tau,ij})\right).$$

The longitudinal outcome was generated as

$$Y_{ij} = \boldsymbol{x}_{ij}^T\boldsymbol{\beta} + S_{\beta i} + \boldsymbol{z}_{ij}^T\boldsymbol{\gamma}_i + b_i^{\beta} + \varepsilon_{ij}.$$

The PVE calibration was modified to include the additional relatedness-induced variation. In particular,

$$c_\tau = \left\{ \frac{\mathrm{PVE}_\tau}{1 - \mathrm{PVE}_\tau} \frac{\mathrm{Var}(\omega + \boldsymbol{b}^\tau)}{\mathrm{Var}(S_\tau^{(0)})} \right\}^{1/2},$$

$$c_\beta = \left\{ \frac{\mathrm{PVE}_\beta}{1 - \mathrm{PVE}_\beta} \frac{\mathrm{Var}\left(\boldsymbol{Z}\boldsymbol{\gamma} + \boldsymbol{\varepsilon} + \boldsymbol{b}^\beta\right)}{\mathrm{Var}(S_\beta^{(0)})} \right\}^{1/2}.$$

All other phenotype-generation settings followed Supplementary Method 9.

### Supplementary Figures

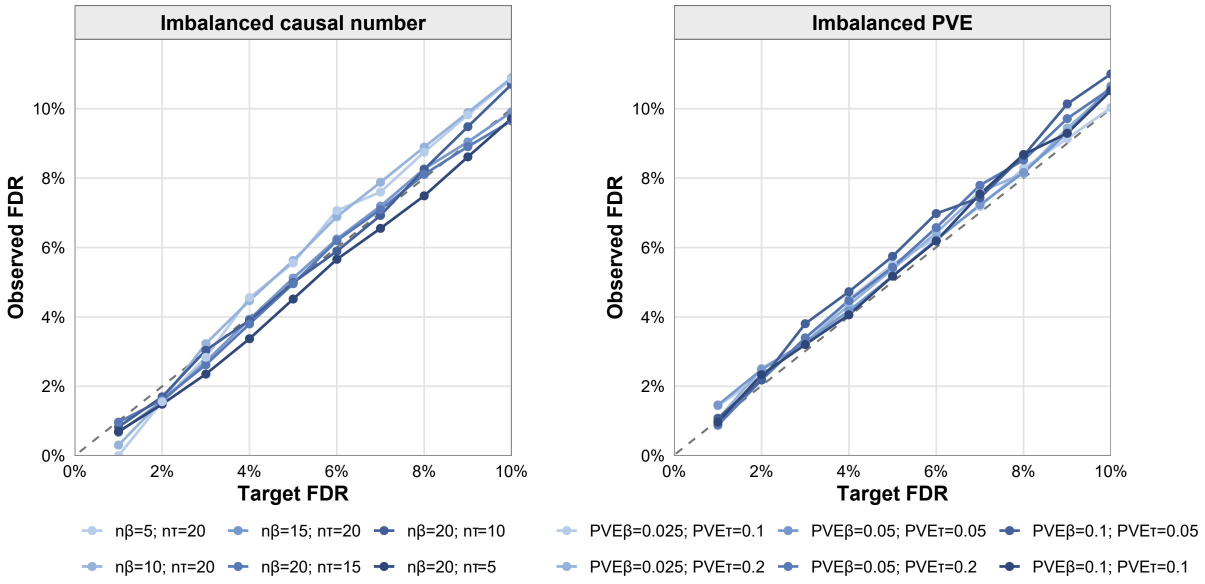

**Figure S1. FDR control under imbalanced genetic architectures.** Observed FDR for LKOM when the two response channels differed in the number of causal variants or channel-specific PVE. The dashed line denotes the target FDR.

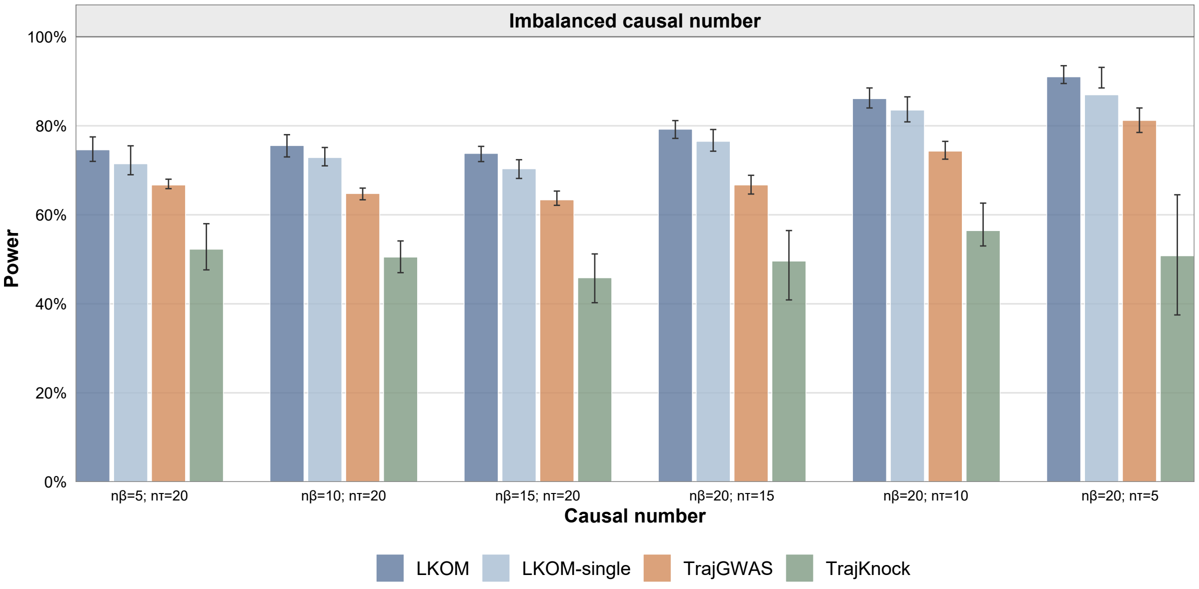

**Figure S2. Power under imbalanced numbers of causal variants.** Channel-wise power when the numbers of causal variants differed between the mean and WS variability channels. Bars show mean power and error bars indicate the interquartile range.

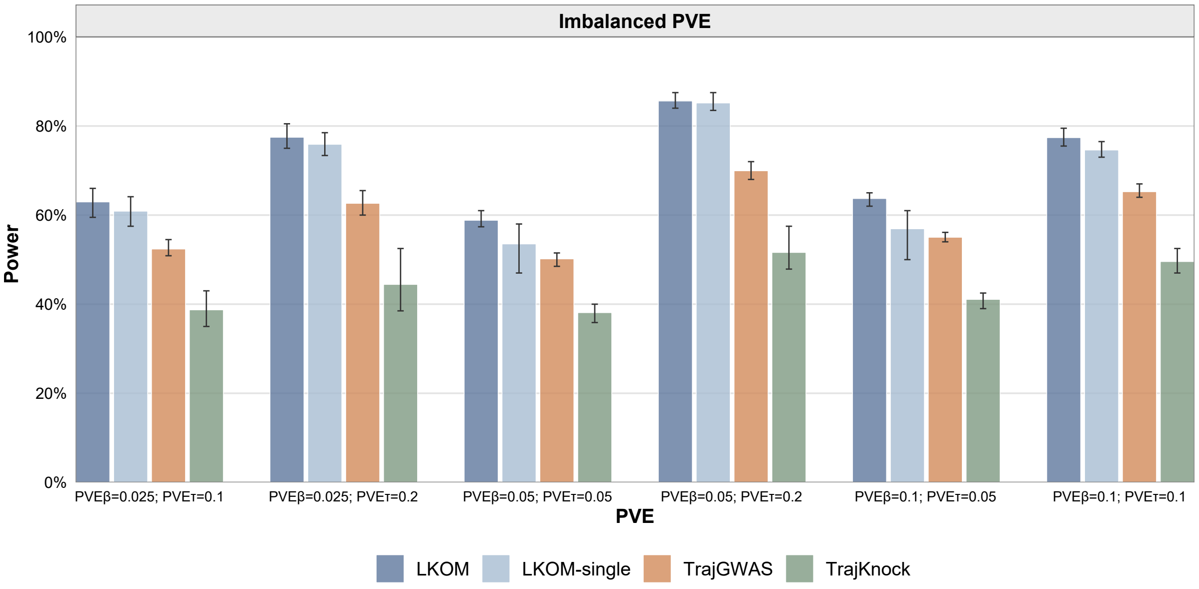

**Figure S3. Power under imbalanced channel-specific PVE.** Channel-wise power of LKOM and the comparison methods under unequal PVE allocations between the mean and WS variability channels. Bars show mean power and error bars indicate the interquartile range.

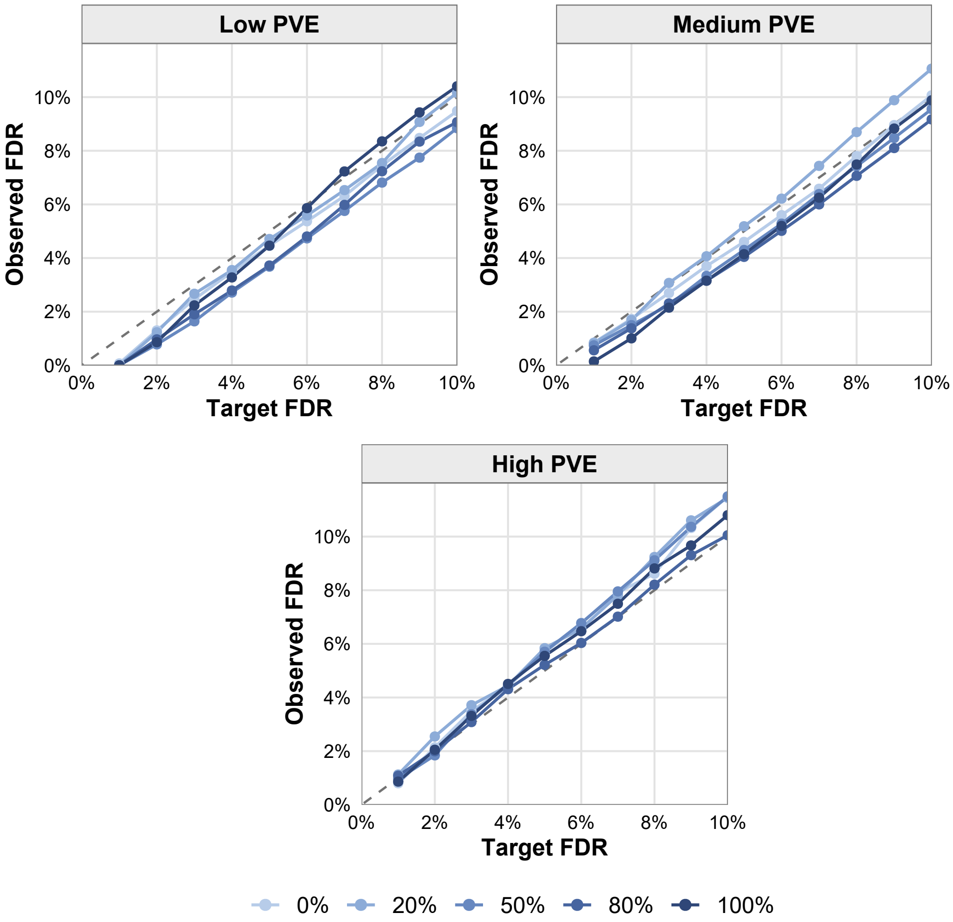

**Figure S4. FDR control across causal-overlap proportions.** Observed FDR of LKOM across different proportions of causal variants shared between the mean and WS variability channels and across PVE levels. The dashed line denotes the target FDR.

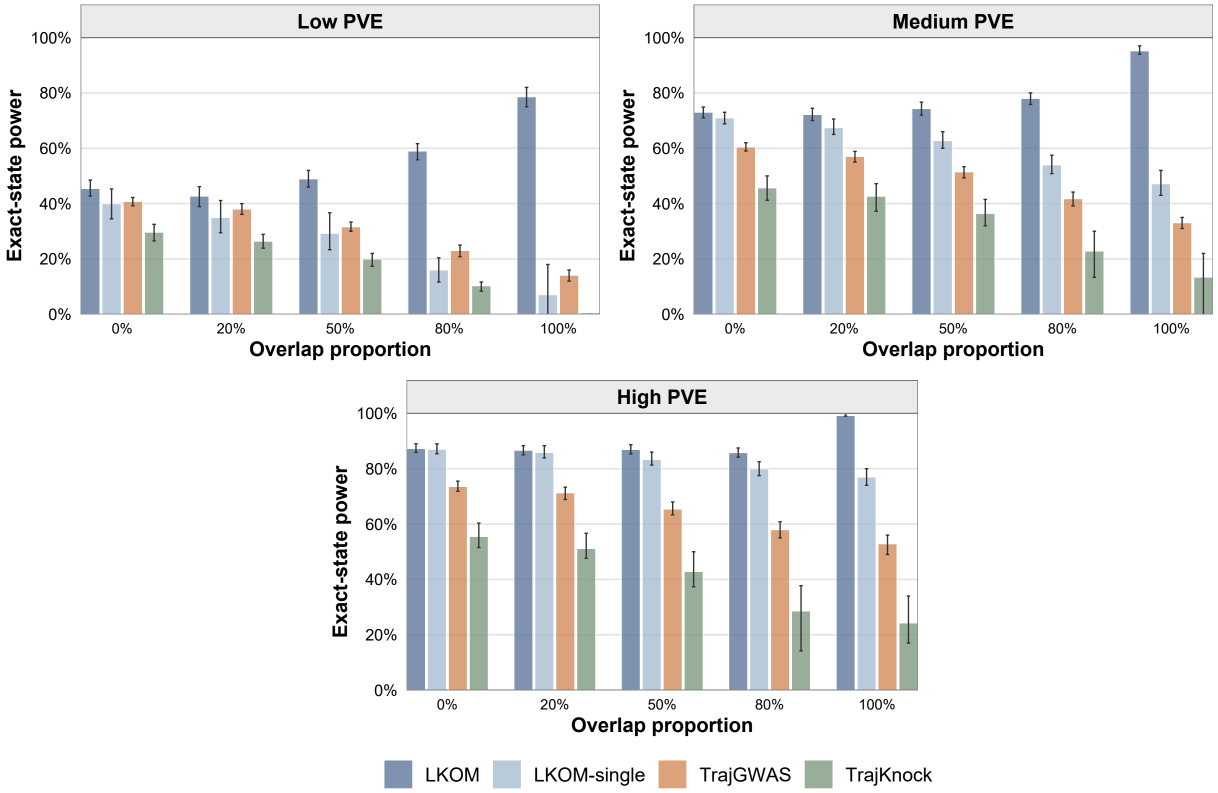

**Figure S5. Exact-state power across causal-overlap proportions.** Exact-state power of LKOM and the comparison methods across causal-overlap proportions and PVE levels. A causal variant was counted only when it was detected and assigned to its correct association state.

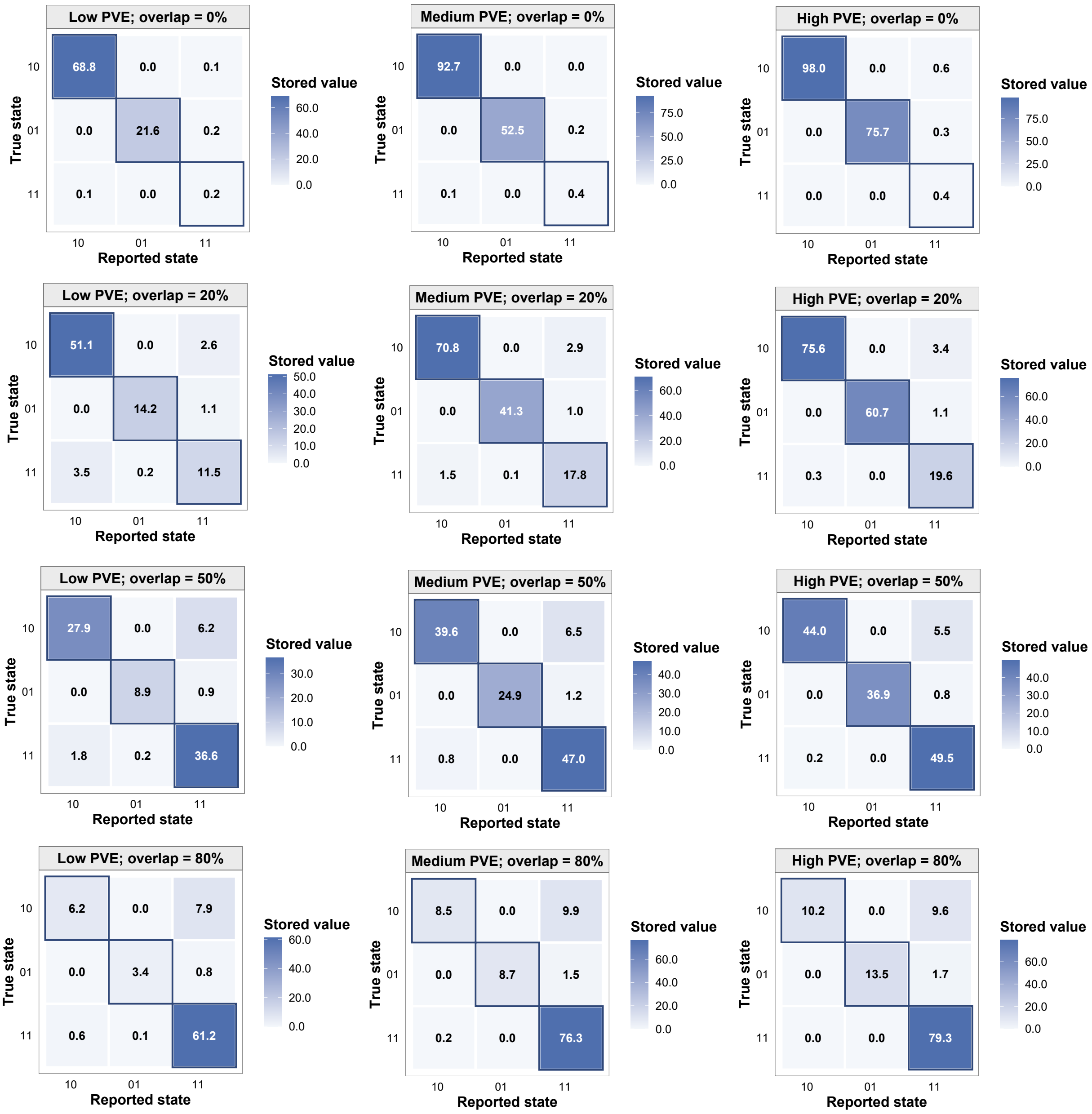

**Figure S6. Association-state classification across causal-overlap proportions.** Correspondence between the true and reported association states 10, 01, and 11 across causal-overlap proportions and PVE levels. Cell entries show the mean numbers of variants assigned to each state across grouped evaluations, with diagonal cells representing correct classification.

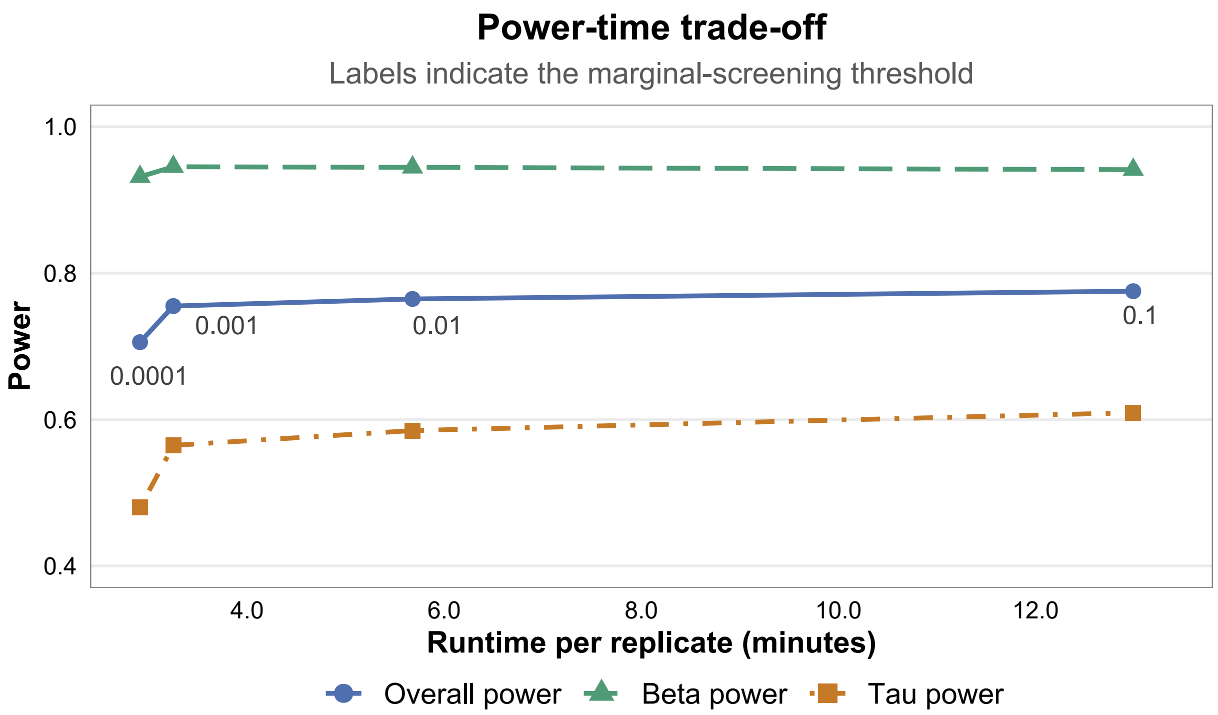

**Figure S7. Power-runtime trade-off across marginal-screening thresholds.** Overall and channel-specific power plotted against runtime per simulation replicate. Point labels indicate the marginal-screening threshold $p_{\text{scr}}$.

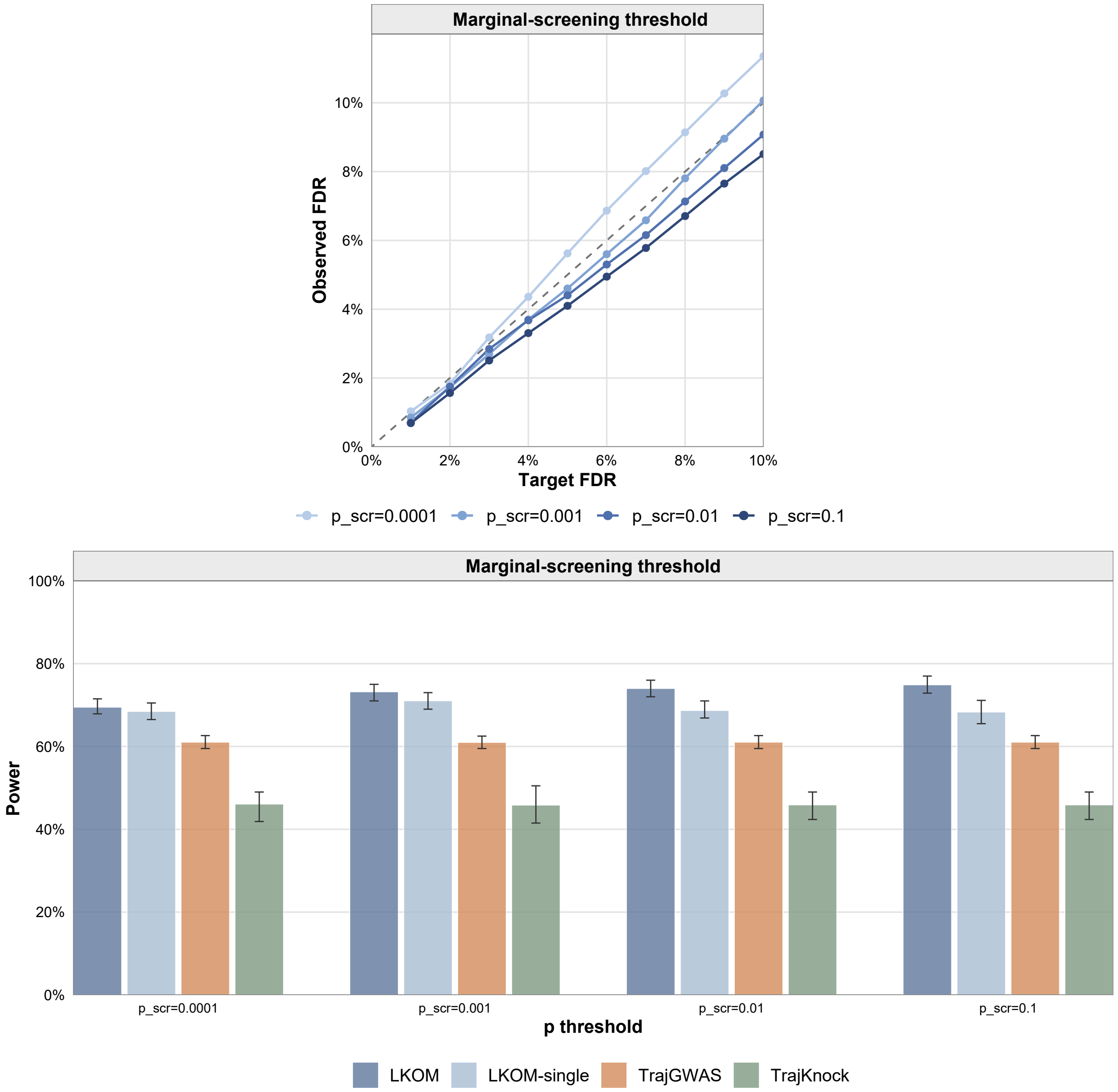

**Figure S8. Effect of the marginal-screening threshold on FDR and power.** Observed FDR of LKOM and channel-wise power of all methods across four marginal-screening thresholds. The dashed line denotes the target FDR; error bars indicate the interquartile range.

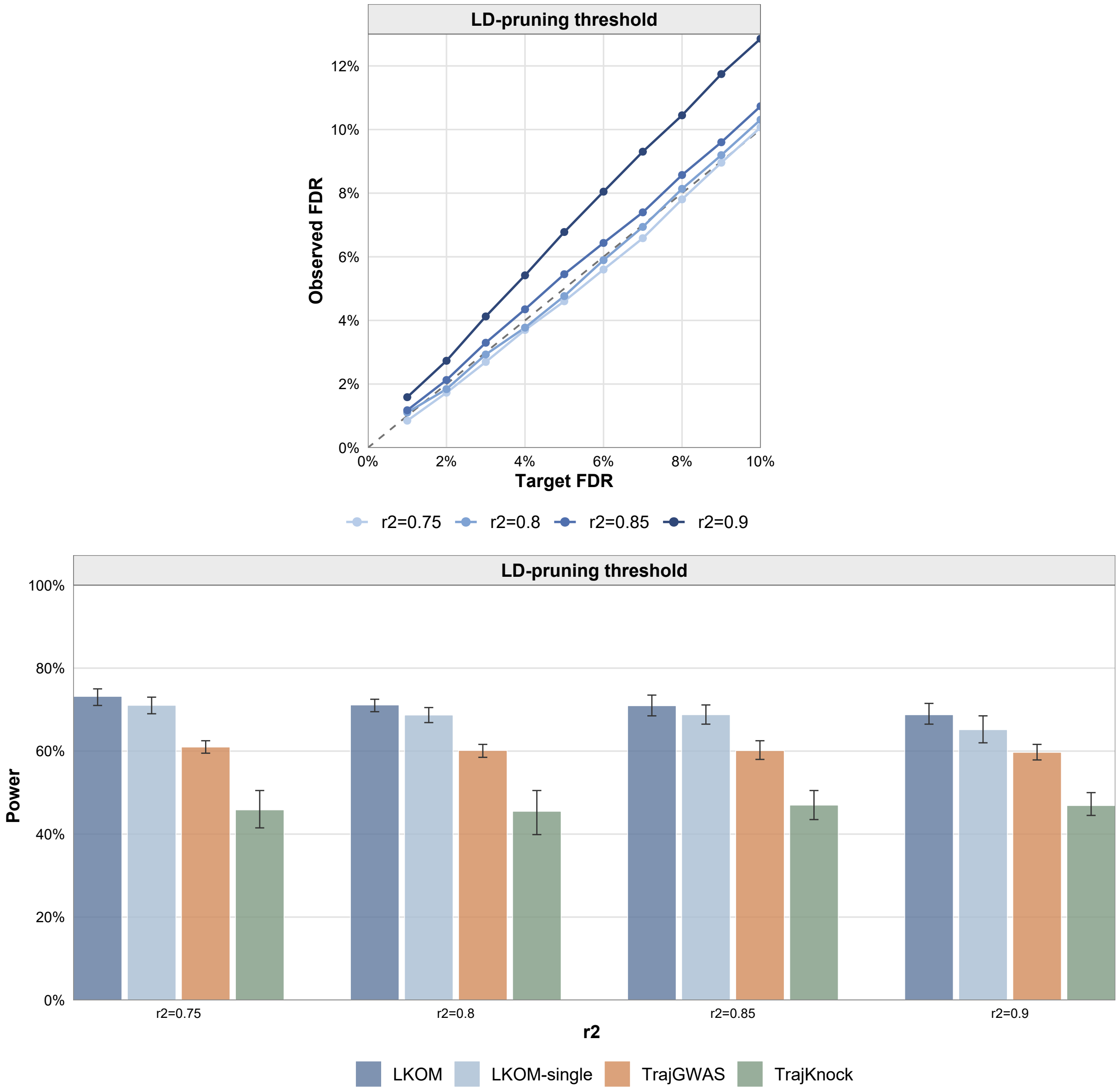

**Figure S9. Effect of the LD-pruning threshold on FDR and power.** Observed FDR of LKOM and channel-wise power of all methods across LD-pruning thresholds $r^2 = 0.75$, 0.80, 0.85, and 0.90. The dashed line denotes the target FDR; error bars indicate the interquartile range.

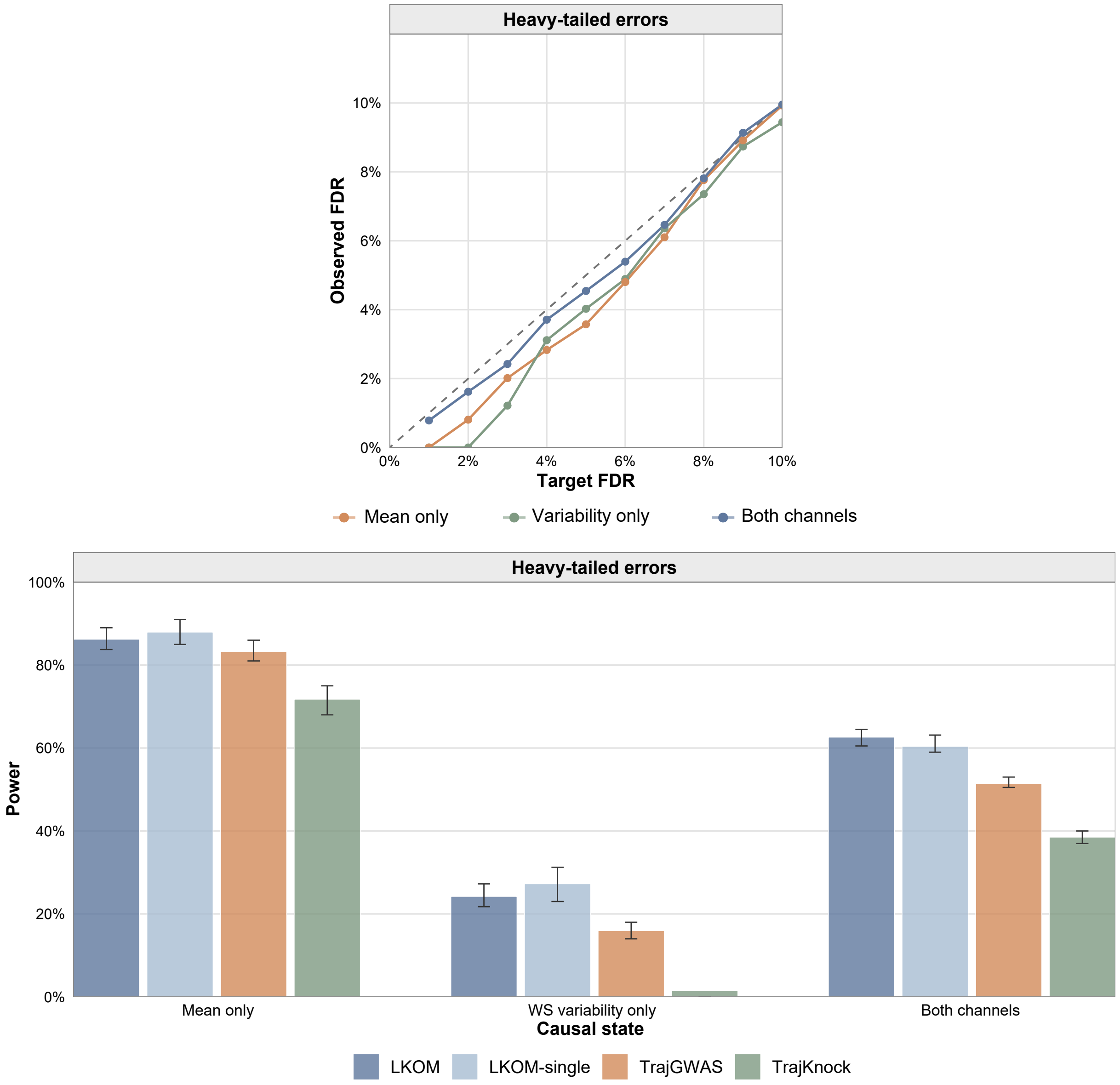

**Figure S10. Performance under heavy-tailed residual errors.** Observed FDR and power under variance-standardized $t_5$ residual errors. The dashed line denotes the target FDR. Bars show mean power for mean-only, WS-variability-only, and dual-channel causal variants; error bars indicate the interquartile range.

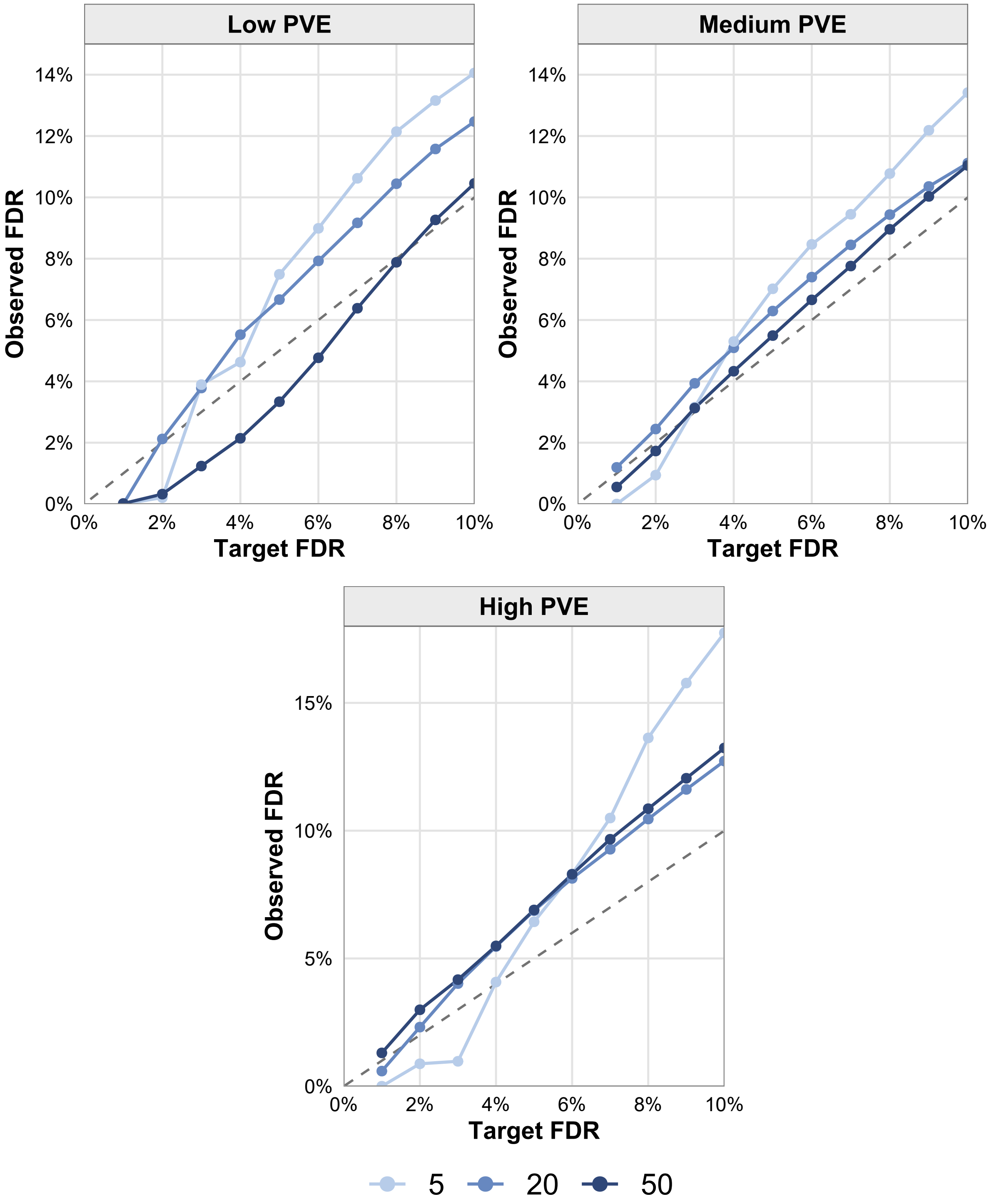

**Figure S11. FDR control without inverse-normal transformation.** Observed FDR of LKOM without inverse-normal transformation across different numbers of causal variants and PVE levels. The dashed line denotes the target FDR.

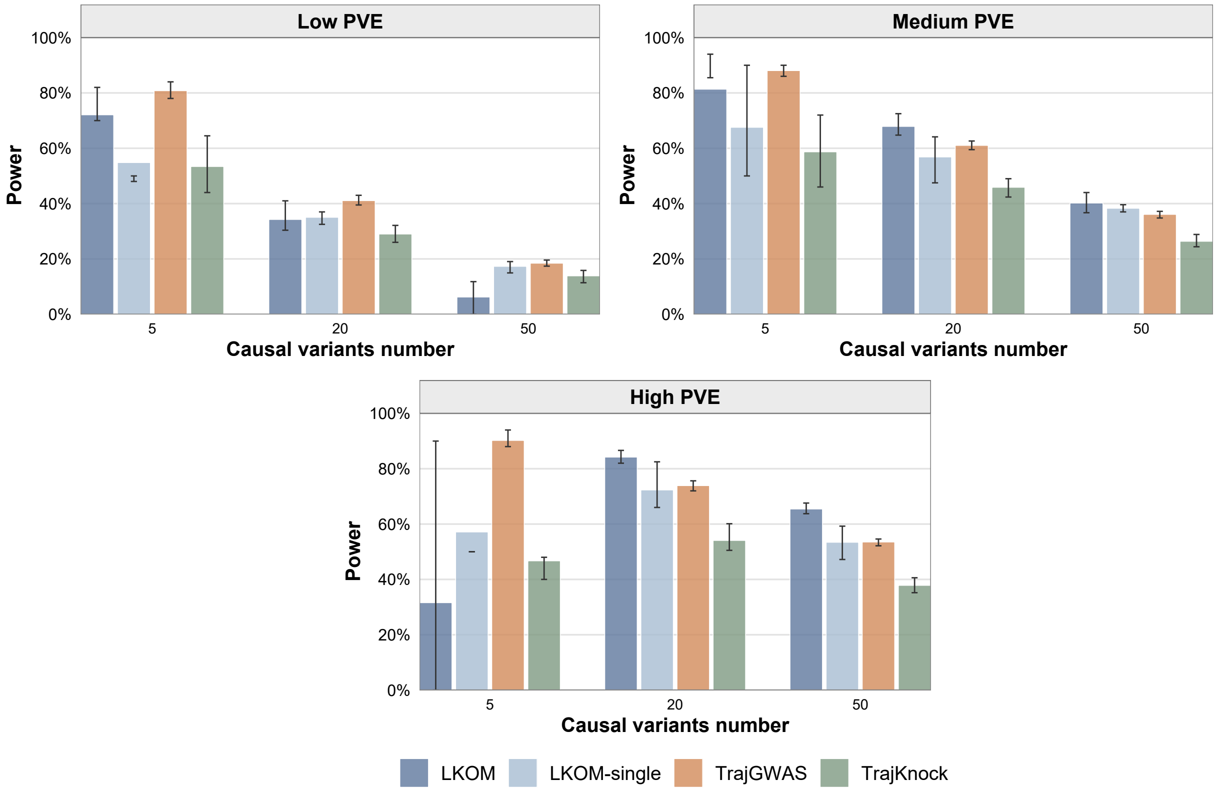

**Figure S12. Power without inverse-normal transformation.** Channel-wise power of LKOM and the comparison methods without inverse-normal transformation across causal-variant numbers and PVE levels. Bars show mean power and error bars indicate the interquartile range.

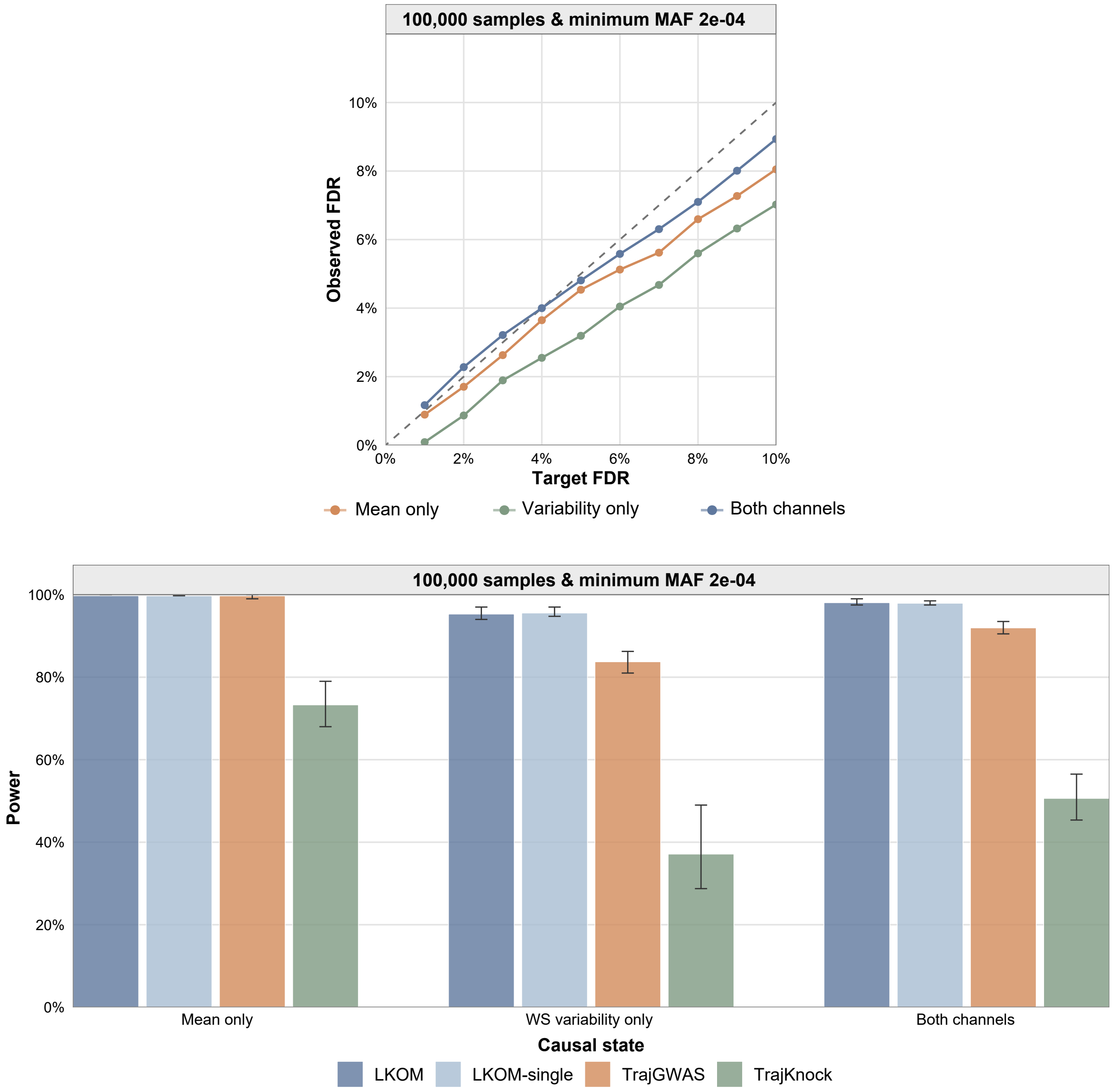

**Figure S13. Performance of LKOM in 100,000 samples.** Observed FDR and power in simulations with 100,000 samples and a minimum MAF of $2 \times 10^{-4}$. The dashed line denotes the target FDR. Bars show mean power for mean-only, WS-variability-only, and dual-channel causal variants; error bars indicate the interquartile range.

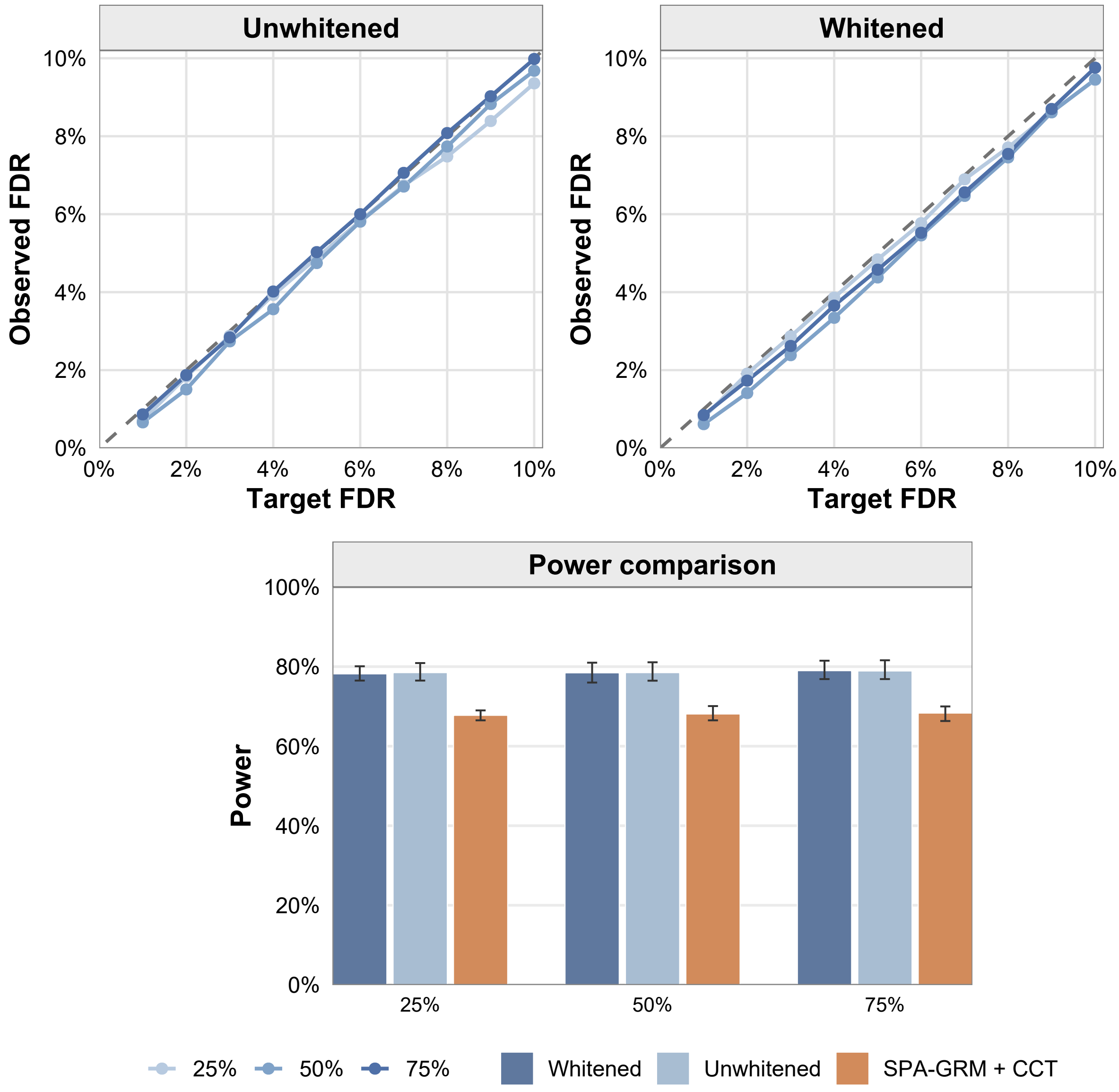

**Figure S14. Performance in UK Biobank-derived cohorts with sparse relatedness.** Observed FDR of LKOM without (top left) and with (top right) GRM-based whitening in cohorts of 10,000 participants, of whom 25%, 50%, or 75% had at least one relative within the third degree in the same cohort. Grey dashed lines indicate equality between observed and target FDR. The bottom panel compares the channel-wise power of whitened LKOM, unwhitened LKOM, and $SPA_{GRM(CCT)}$ at a target FDR of 5%. Bars show means and error bars indicate the interquartile range across 500 grouped evaluations.

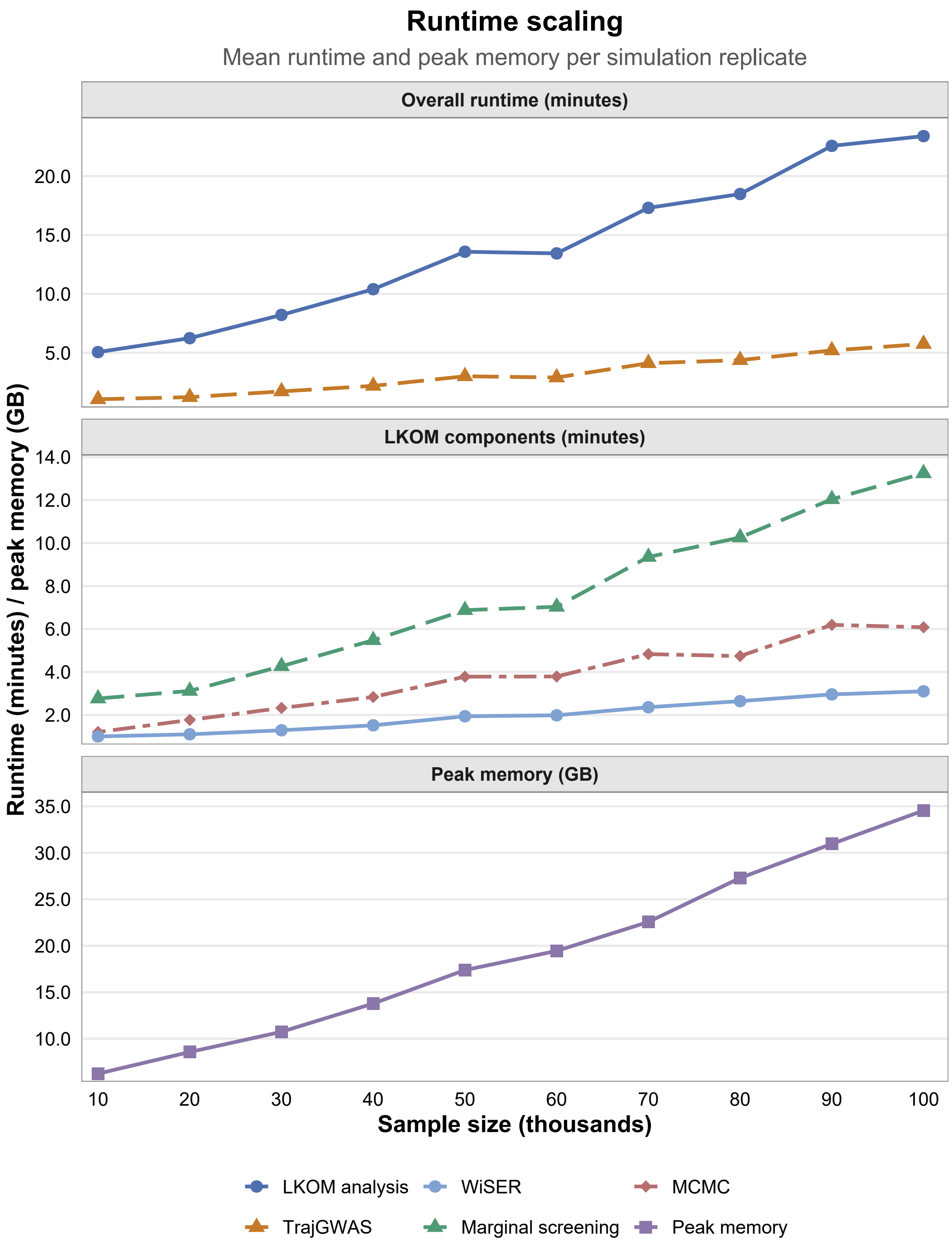

**Figure S15. Runtime and memory scaling with sample size.** Mean runtime and peak memory per simulation replicate across sample sizes from 10,000 to 100,000. The upper panel shows the overall runtime of LKOM and TrajGWAS, the middle panel shows the runtime of the WiSER, marginal-screening, and MCMC components of LKOM, and the lower panel shows peak memory usage. LKOM runtime excludes knockoff construction.

### Supplementary Tables

**Table S1.** Summary of the simulation scenario grid. Only parameters varied across simulation experiments are shown; the data-generation and evaluation procedures are described in the Methods.

| Analysis | Varied parameter(s) | Scenario values |
|---|---|---|
| Association states | State and signal strength | 00 under the global null; 10, 01, and 11 under $$(\mathrm{PVE}_\beta, \mathrm{PVE}_\tau) \in \{(0.025, 0.05), (0.05, 0.10), (0.10, 0.20)\},$$ with the PVE of an inactive channel set to zero. |
| Causal-number analysis | Number of causal variants and PVE | $$\lvert\mathcal{C}_\beta\rvert = \lvert\mathcal{C}_\tau\rvert \in \{5, 20, 50\}$$ crossed with the three PVE pairs, giving nine scenarios. |
| Overlap analysis | Cross-channel overlap and PVE | $$\omega \in \{0, 0.2, 0.5, 0.8, 1\}$$ crossed with the three PVE pairs, giving 15 scenarios. Twenty causal variants were included in each channel. |
| Asymmetric causal architecture | Channel-specific causal numbers | $$(\lvert\mathcal{C}_\beta\rvert, \lvert\mathcal{C}_\tau\rvert) \in \{(5, 20), (10, 20), (15, 20), (20, 5), (20, 10), (20, 15)\}.$$ |
| Unequal signal strength | Channel-specific PVE | $$(\mathrm{PVE}_\beta, \mathrm{PVE}_\tau) \in \{(0.025, 0.10), (0.025, 0.20), (0.05, 0.05), (0.05, 0.20), (0.10, 0.05), (0.10, 0.10)\}.$$ |
| Simulated family structures | Family size | 5,000 unrelated individuals combined with 1,250 four-member families, 500 ten-member families, or 250 twenty-member families. |
| UK Biobank kinship cohorts | Proportion of related participants | 25%, 50%, or 75% of participants had at least one third-degree-or-closer relative in the analysis cohort. |
| Related-sample analysis | GRM-based whitening | LKOM with and without GRM-based whitening. |
| Transformation sensitivity | Residual inverse normal transformation | Applied versus omitted across the balanced causal-number–PVE scenarios. |
| Posterior-computation sensitivity | Inference procedure | Gibbs posterior sampling versus the deterministic conditional Bayes factor procedure. |
| LD-pruning sensitivity | LD-pruning threshold | $r^2 \in \{0.75, 0.80, 0.85, 0.90\}$. |
| Screening sensitivity | Marginal screening threshold | $p_{\mathrm{scr}} \in \{0.0001, 0.001, 0.01, 0.1\}$. |
| Distributional sensitivity | Conditional residual distribution | Normally distributed versus variance-standardized $t_5$ innovations. |
| Follow-up sensitivity | Number of measurements | Uniformly sampled from $\{6, \ldots, 15\}$ versus $\{2, \ldots, 8\}$. |
| Scalability analysis | Sample size and minimum MAF | $n$ = 10,000 with minimum MAF $1.25 \times 10^{-3}$ versus $n$ = 100,000 with minimum MAF $2 \times 10^{-4}$. |

Unless otherwise specified, the default setting used 10,000 participants, 20 causal variants per channel, $(\mathrm{PVE}_\beta, \mathrm{PVE}_\tau) = (0.05, 0.10)$, no imposed cross-channel overlap, $r^2 = 0.75$, $p_{\mathrm{scr}} = 0.001$, normally distributed conditional residuals, and 6–15 longitudinal measurements per participant. Each sensitivity analysis varied one component while keeping the remaining settings fixed.

**Table S5.** Mean numbers of detected causal variants according to their true association states. Each state contained 10 causal variants per simulation replicate.

| Target $q$ | PVE | True state | LKOM | LKOM-single | TrajGWAS | TrajKnock |
|---|---|---|---|---|---|---|
| 0.05 | Low | 10 ($\beta$ only) | 6.84 | 6.88 | 6.73 | 5.78 |
| | | 01 ($\tau$ only) | 2.04 | 0.95 | 1.45 | 0.12 |
| | | 11 (both) | 7.65 | 3.84 | 4.09 | 2.90 |
| | Medium | 10 ($\beta$ only) | 9.24 | 9.31 | 8.70 | 7.39 |
| | | 01 ($\tau$ only) | 5.26 | 4.72 | 3.43 | 1.75 |
| | | 11 (both) | 9.52 | 7.11 | 6.03 | 4.45 |
| | High | 10 ($\beta$ only) | 9.87 | 9.90 | 9.62 | 7.86 |
| | | 01 ($\tau$ only) | 7.63 | 7.48 | 5.20 | 2.93 |
| | | 11 (both) | 9.93 | 8.81 | 7.41 | 5.13 |
| 0.10 | Low | 10 ($\beta$ only) | 7.39 | 7.43 | 6.97 | 6.67 |
| | | 01 ($\tau$ only) | 2.47 | 2.03 | 1.63 | 0.65 |
| | | 11 (both) | 8.01 | 4.75 | 4.28 | 3.57 |
| | Medium | 10 ($\beta$ only) | 9.40 | 9.46 | 8.82 | 8.35 |
| | | 01 ($\tau$ only) | 5.62 | 5.28 | 3.66 | 2.68 |
| | | 11 (both) | 9.59 | 7.49 | 6.22 | 5.47 |
| | High | 10 ($\beta$ only) | 9.89 | 9.91 | 9.67 | 8.89 |
| | | 01 ($\tau$ only) | 7.69 | 7.63 | 5.43 | 4.07 |
| | | 11 (both) | 9.94 | 8.87 | 7.56 | 6.31 |

**Table S7.** Mean numbers of causal and detected causal variants across MAF strata.

| Target $q$ | PVE | MAF stratum | Causal | LKOM | LKOM-single | TrajGWAS |
|---|---|---|---|---|---|---|
| 0.05 | Low | Rare | 12.77 | 7.07 | 4.49 | 3.53 |
| | | Low-frequency | 3.81 | 2.94 | 1.95 | 2.29 |
| | | Common | 3.43 | 2.08 | 1.33 | 2.33 |
| | Medium | Rare | 12.77 | 10.24 | 8.32 | 6.03 |
| | | Low-frequency | 3.83 | 3.61 | 3.31 | 3.12 |
| | | Common | 3.40 | 2.86 | 2.43 | 2.95 |
| | High | Rare | 12.77 | 11.67 | 10.66 | 8.13 |
| | | Low-frequency | 3.80 | 3.76 | 3.72 | 3.46 |
| | | Common | 3.43 | 3.25 | 3.13 | 3.23 |
| 0.10 | Low | Rare | 12.77 | 7.56 | 5.35 | 3.78 |
| | | Low-frequency | 3.81 | 3.09 | 2.45 | 2.37 |
| | | Common | 3.43 | 2.26 | 1.69 | 2.39 |
| | Medium | Rare | 12.77 | 10.41 | 8.71 | 6.30 |
| | | Low-frequency | 3.83 | 3.65 | 3.45 | 3.18 |
| | | Common | 3.40 | 2.99 | 2.66 | 2.97 |
| | High | Rare | 12.77 | 11.68 | 10.70 | 8.36 |
| | | Low-frequency | 3.80 | 3.76 | 3.73 | 3.49 |
| | | Common | 3.43 | 3.28 | 3.21 | 3.25 |

Rare, $0.00125 \leq \text{MAF} < 0.01$; low-frequency, $0.01 \leq \text{MAF} < 0.05$; common, $0.05 \leq \text{MAF} \leq 0.5$.
